# A transcriptome analysis of *OsNAC02* Ko-mutant during vegetative endosperm development

**DOI:** 10.1101/2023.04.10.536330

**Authors:** Mei Yan, Guiai Jiao, Gaoneng Shao, Ying Chen, Maodi Zhu, Lingwei Yang, Lihong Xie, Peisong Hu, Shaoqing Tang

## Abstract

The Ko-*Osnac02* mutant matured earlier than the wild type (WT) in 15 days obviously. The result showed that the *Ko-Osnac02* chalky with white-core and white-belly mature endosperm. The seed filling rate plays a major role in determining the yield and quality of rice (Oryza sativa L. versus japonica). Interestingly, in the Ko-*Osnac02* seeds, higher amylose content (AC) was observed in vegetative endosperm (5DAP), while lower amylose content (AC) and higher soluble sugar content were observed in the mature endosperm. RNA-Seq analysis of N2/N3 seed (3DAPs) and the WT revealed that among the top differentially expressed genes (DEGs), the OsBAM2 (LOC_Os10g32810) expressed significantly high which involved in starch degradation. In addition, seven genes were expressed at a lower-level in pro-pro interactions inducing the chalky endosperm formation in N2 mutant seeds (3 DAP), which could be verified by heatmap diagrams based on DEGs of N2 versus WT. The cell cycler controlling the Tubulin genes downregulated their expression together with the MCM family genes MCM4 ( ↓ ), MCM7 ( ↑ ), which may cause white-core in the early endosperm development. In conclusion, the developing period shorten significantly in the Ko-*Osnac02* mutants may cause the chalkiness in seeds during the early endosperm development.

**ONE SENTENCE SUMMARY:** The gene *OsNAC02* which controls a great genetic co-network for cell cycle regulation in early development, and Ko-*Osnac02* mutant shows prematurity and white-core in endosperm.

## Introduction

Rice (*Oryza sativa* L.) is one of the most important cereal crops in the world, it serves as the main dish for more than half of the world’s population, and it is a major food for Asians. Rice is mainly planted in Asia and Africa. It is estimated that rice production need increasing by 30% in 2030, due to the growing population and increasing rice consumption in China[1]. The extreme climate is the crucial limiting factor for global production of cereals such as rice, maize and wheat [2]. Specifically, climate change accompanied with hydration, water flooding, disease stress caused yield losses worldwide in 2022. A growing environment with hydration resistance with salt stress, and flood resilience with wounding tolerance has become an urgent goal for the improvement of crops for the next generation. Filling rate is a very important factor for yield and seed development period determination [3, 4].

In this study, the grain filling rate was accelerated in the Ko-*Osnac02* mutant compared with the wild type (WT), which caused the mediant to be chalky in the endosperm of mature seeds (125 DAG, days after germination). This induced the early maturity which not only affected the yield, but also decreased the quality simultaneously. Thus, RNA-seq analysis of the Ko-mutant seeds (3 DAP, days after pollination) was primarily performed and online bioinformatics were used for the motif detection in promoter regions. Meanwhile, the multi-regulation of OsNAC02/OsNAC06 at the mRNA-protein level was revealed by multi-omics such as pro-pro String and Mapman frack, in the Sucrose and Starch metabolic pathway, and the environmental stress response at protein-level could continuously display the mechanisms of the premature and chalky phenotype.

The bioinformatics pro-pro String analysis revealed that OsNAC02 could regulated more than five effective TFs, which has been verified by H-1-Y including HSF, NAC and MYB factors, meaning the abiotic resistance of the *OsNAC02* Ko-mutant changes according to the pathway activated by these TFs. Heat shock promoters are characteristicized by the conserved palindromic elements with the consensus motif “-nGAAnnTTCn-” among eukaryotes[5, 6]. However, the cis-element ABRE is also enriched in the promoter region of HSF in barley[7], and the “ACGT” motifs, such as the G-boxes and ABRE motifs, are also enrichment in the promoter of these genes, and they could respond to abiotic stress and heat stress[8–12]. In a previous study, except HSF, the TFs including NAC, AP2/ERF, WRKY, MYB, bHLH, MADS and C2H2 predominantly responded to heat stress [13–15]. All of these TFs are directly regulated by OsNAC02 and have the ABRE/G-box in their promoters, which means that they could not only respond to heat shock, but they could be induced by other abiotic stress. NAC(NAM-ATAF-CUC) was a plant-specific transcription factor which has a large family and is distributed widely in plants (S3 Fig.), but it has a similar function in homologous plants [16, 17]. The rice NAC-domain proteins are highly conserved in different plants, and they all have similar abiotic stress responses and tolerance to harsh environments [3, 18–25].

*OsNAC02* (LOC_Os02g38130, Os02g0594800) was in the upstream of cell cycle control. It is located in the nucleus where it regulates the development of various organs in rice, such as the seed, flower, SAM and root of seedlings. The vegetative and reproductive development processes in *OsNAC02* Ko-mutant may suppress starch synthesis by the Starch and Sucrose pathways, which are implicated to change the endosperm development as a result.

In this study, the *OsNAC02* Ko-mutants were created on the nipponbare background. Compared with the WT, the homeostasis of starch metabolism in the *OsNAC02* Ko-mutant induced a premature and chalky phenotype in the mutant seeds.

In full-filled seeds, *OsNAC02* knock-out will affect the morphological characters of the Ko-mutant including a decrease in the quality of endosperm and an acceleration in seed development period obviously. Because OsNAC02 could interact with more than five TF proteins simultaneously, and its mRNA is unique in the upstream, which means that they all control the cellular process through a specific mechanism, and the effective expression change of these TFs may induce energy imbalance and activation of the phytohormone pathways. From these results, we found that a small change in *OsNAC02* expression may indirectly affect the important pathways related to cell division.

## Materials and Methods

### The construction of Ko-*Osnac02* and KO-*Osnac06* mutants

The KO-mutants are constructed by the Crispr-Cas9-Vector. Both N2 and N3 *mut* are multiple mutants, and we obtained homozygote transplants from T1 and T2 generations in 2020 and 2021 separately. The mutant creation vectors are constructed by co-infusion to the Ku-Cas9pl-Ko-vector (Kan^+^), the tags designed on the CDS of gene sequences with **NGG/NAG** as PAM-seq (Fig. 1A and 1B). The two Oligo diploymers are detected in T0 generation by the primers designed on mutant sites, with the target region designed to be −10 bp upstream of ‘ATG’. The tags in the Crispr-Cas system are designed by software online (http://www.biogle.cn/index/excrispr) or (http://crispr.dbcls.jp/), and the sequence input online are both CDS of genes, the tags are selected from output result list with consecutive sequence in CDS regions by alignment with gene DNA sequence.

**Fig 1.**
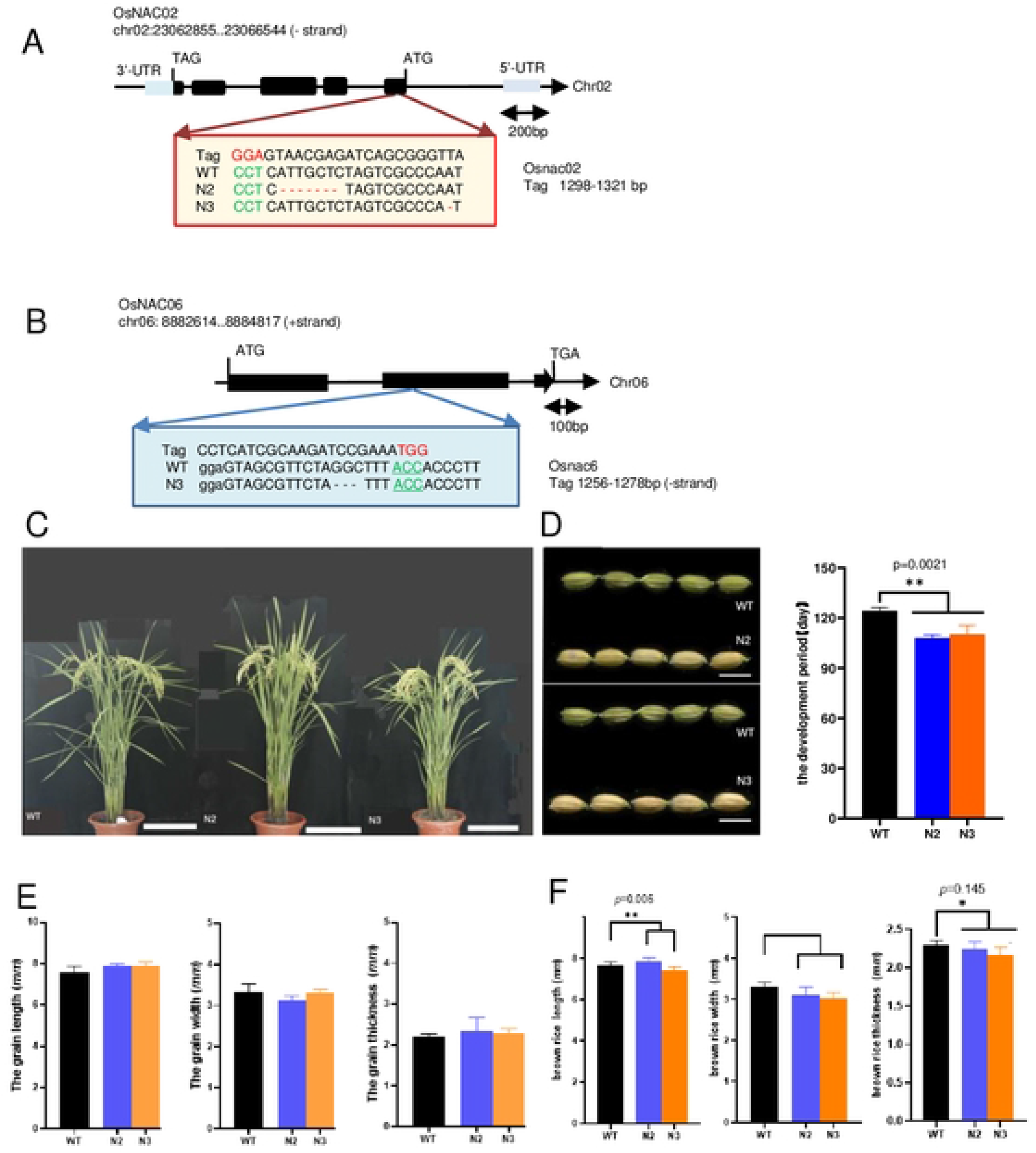
The KO-OsNAC02/0sNAC06 mutant (T2) construction and botanic characters detection. Two mutants named N2, N3 are planted in field (Fuyang, Hangzhou in 2021), andthe homozygous was harvested in the autumn. The control is wild type of nipponbare. The genetic detection of all homozygous mutants was performed by PCR sequencing. The N2 has two mutant sites in OsNAC02 CDS regions, N3 has one mutant site in OsNAC02 and OsNAC06 CDS regions respectively (**Figure S1**). **(A)** N2 mutant construction. The KO-Osnac02 mutant created by Osnac02 - tag2 has 7 bp-deletion in N2 mut and KO-Osnac02 mutant has 1bp-deletion in N3 mut, respectively. **(B)** N3 mutant construction. The KO-Osnac06 mutant created by Osnac06 - tag2 has 3 bp-deletion in N3 mut. **(C)** The N2/N3 mut plants (95DAG) are selected for botanic characters detection(**Figure S2**). n= 17.5 cm. **(D)** The premature phenotype in N2/N3 mut (95DAG) and the development period calculation of the mutants (two-way ANOVA for mixed model, *, p < 0.05 and **, p < 0.01, n= 6 mm).The mutant matured earlier than WT as a result. For the detection of premature phenotypes in N2 andN3 mut, the seed shape characters such as length, width, and thickness are collected in **Fig E** and **Fig F**. **(E)** The shape phenotypes detection of N2/N3 mut seeds (95 DAG), and there are no difference between mut vs WT. **(F)** The shape phenotypes of full-mature seeds (T_2_) in N2/N3 mut (125 DAG) are detected, which were harvested in 2021. The full-filledseeds (125 DAG) of WT is larger than mutants on the whole.

In order to detect the function of OsNAC02, the mutant transplants of T0 generation were obtained and relative genes expressions are detected by qRT-PCR (**Fig.** 2A and 2D).

**Fig 2.**
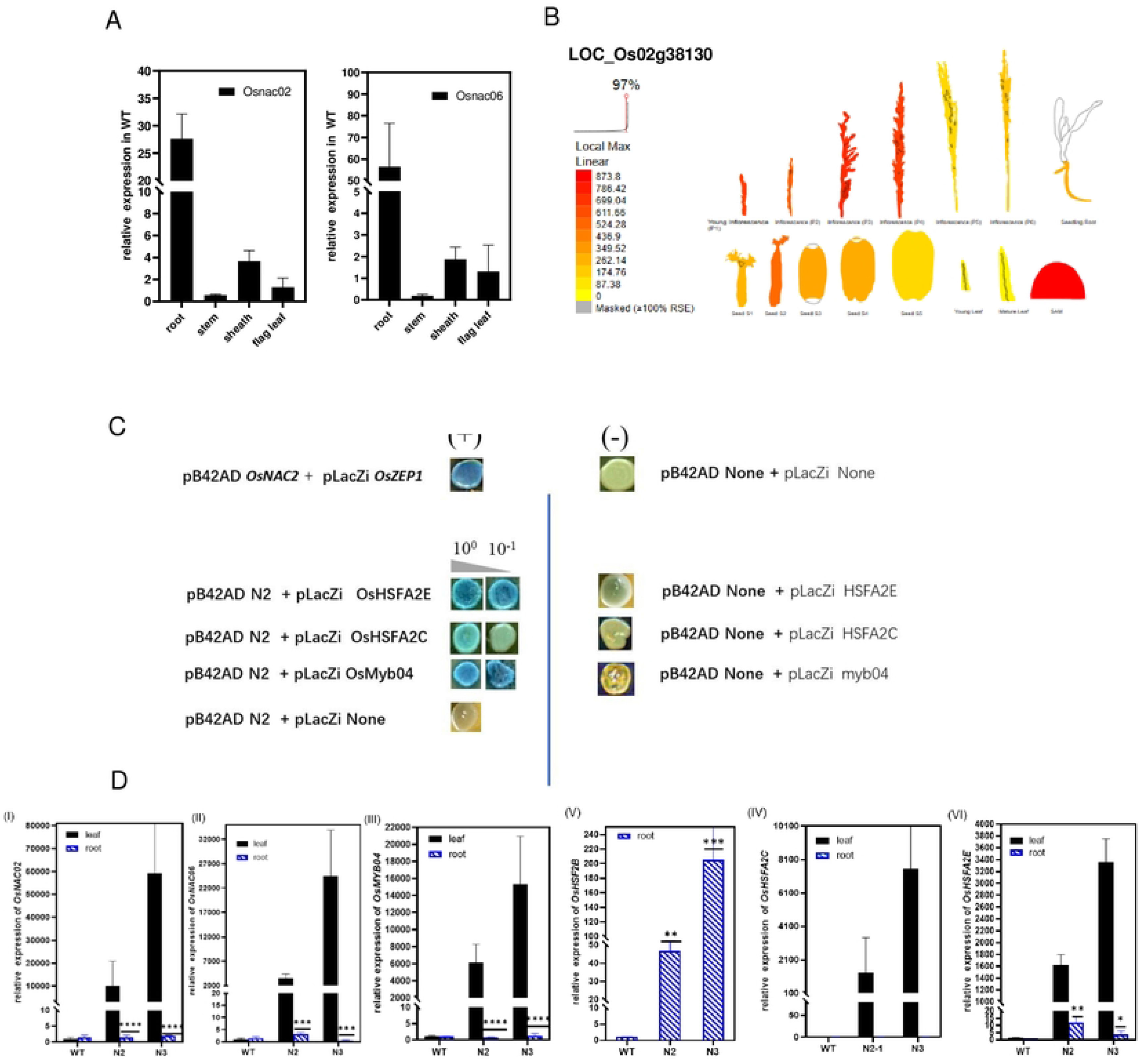
The functional analysis of OsNAC02 and its co-network construction. **(A)** The OsNAC02 and OsNAC06 distinct expressions in different tissues verilied by qRT-PCR. Both of them expressed highly in root, as this organ is most vegetative parts of WT seedling. The ubiquitin(LOC_Os03g13170.1, Os03g0234200Jwas used as the reference which expressed in the whole plant. **(B)** The distinct tissue expressions of OsNAC02 (LOC_Os02g38130,Os02g0594800) on the Plant eFP Viewer (http://bar.utoronto.ca/eplant_rice/). and this result is in accordance with **Fig A**. **(C)** Yeast-one-hybrid assays show the OsNAC02 interact with the promoter fragments of OsHSFA2C, OsHSFA2E, OsMYB04. The positive control in bluecolonies was constructed according to the reference (Mao et al., 2017). There are two cultures in this experiment used as the selection medium: SD and SD /-Ura /-Trp. The X-gal culture is SD /-Ura /-Trp + DMF + X-gal with the content of 10^-1^ and 10^-2^ for positive screen. **(D)** The qRT-PCR results represent the expressions of OsNAC02 regulated genes in the leaf and root of mutants vs WT, including: OsNAC02, OsNAC06, 0sMYB04, 0sHSF2B, OsHSFA2C, OsHSFA2E. The ubiquitin (LOC_Os03g13170.1) is the reference gene (student t·test, ·, p < 0.05 and”, p <0.01 J, and the motif locations are listed in the results (**Figure S5A**).

### The yield characters detection and growth conditions

The Nipponbare (*Oryza sativa* L. vs *japonica*) is used as the background of Crispr-Cas9 mutants including the N2 *mut* (*Osnac02*), N3*mut* (*Osnac02/Osnac06*). All the materials were harvested in 2021. The shape phenotypes including the length, width, thickness of grains treated with dehydration in the lab was detected on the seeds from the generation T2 (95 DAG) in the summer of 2021 and T1 (125 DAG) in the winter of 2020.

All the materials are grown in the fields under the normal conditions. The transplants of T0 generation were planted in Lingshui (Hainan island Province,18.5°110’E) during the winter in 2020, where we got T1 generation of mutant. Then transplants of T1 generation are planted in Fuyang (Hangzhou, Zhejiang Province, 30°120’E) during the summer of 2021.We got the T2 generation in the autumn, and the yield characters of mutants are collected in 2021.The important botanic and agronomic characters of the T2 mutants (95 DAG) are detected as follows: the plant height, tiller number, the primary branches and the available panicles number per plants in the field (FuYang, HangZhou, 2021)

All the materials were harvested in 2021. After drying in the air for 60 days, the yield characters are detected including number of seeds per panicle, yield per plant, the weight of 1000-grains per plant. In addition, five randomly selected panicles per plant are used for yield characters detection, such as panicle length, numbers of panicles per plant and number of grains per panicle (S1 **Fig.**).

The manufacture’s peeling caryopsis from seed in a small milled machine, and selected brown rice from a random group of five panicles per plant for the evaluation of yield characters, with the index results from automatic seed testing analyzer with 1000-grain weight meter (Model Type: Wanshen SC-G seed testing instrument). The experimental materials are as follows: N2 *mut* (*Osnac02*), N3 *mut* (*Osnac02/Osnac06*). Besides, the WT (Nipponbare) is used as the control.

### The homologous analysis of OsNAC02 and OsNAC06 for the OsNAM proteins identification

Homologs of *OsNAC02* and *OsNAC06* were identified by BLASTP search settings in the GenBank, these sequences were extracted from NCBI (Uniprot database). These amino acid sequences of 9 predicted proteins in plant mainly belong to rice, maize and Arabidopsis (http://www.ncbi.nlm.nih.gov).

The cluster analysis makes a powerful proof of function that the structure of the N-terminus is conserved while the C-terminus changed variably. The homologous analysis for NAC protein sequences similarity is through using the DNAMAN 7.0 (S4A **Fig.**). The phylogenetic tree of NAC family in rice was constructed in Neibor-Joining cluster by using the MEGA 7.0 (S3 **Fig.**).

### The motif locations in promoter of TFs regulated by OsNAC02

In the OsNAC02 co-network, TFs motifs with ABRE/ G-box having abiotic resistance were screened on promoters of OsMYB4, OsHSFA2c and OsHSFA2e in affyPLM detection, which was used for quality control and producing the genes expression matrixes. (https://www.bioconductor.org/packages/release/bioc/html/affyPLM.html). All the results are listed in Figure S5A. All the motif locations on the promoters of TFs are detected by the PlantCARE online (http://bioinformatics.psb.ugent.be/webtools/plantcare), and the mimic graph of motifs location model was constructed by the software IBS 1.0 (S5B Fig.).

### Cellular observation

The starch granules observation in the central part of transverse section by scanning electron microscopy (SEM). For the observation of mature endosperm starch granules, the cross-sections of mature caryopses were sputter-coated with gold and the semi-thin sections of the opaque endosperm were observed by scanning electron microscope (SEM). The preparation of grain samples was directly dehydrated to a critical point with liquid CO_2_, mounted on SEM stubs and then coated with the gold palladium. The mounted specimens are observed and photographed under the SEM.

The samples of endosperm development observation were fixed with formaldehyde-acetic acid-eth-anol fixative solution and stored at 4 °C, then dehydrated by a gradient ethanol series and infiltration with xylene for embedding in resin for the preparation. The histological analysis in hand-cut sections of filling endosperm of 3 DAP, 5 DAP, 8 DAP, 10 DAP were sliced in 2.0 um-5.0 um-thick sections, and the cellular observation of endosperm development stained with Lugol’s iodine: 0.02% iodine (I_2_) and 0.2% potassium iodide (I_2_-KI), the sections were then observed and photographed under a bright-field microscope (Olympus). All the samples pre-treatment was as follows: the filling samples in different stage were sectioned transversally into semi-thin-slides, then immersed in staining liquid for 30s, then dry the slide on the 37°C.

On the whole, the cell walls and starch granules observation of the endosperm are spanned by the TEM, and the volume, shape and numbers of cells are observed in a same scope area. Meanwhile, the endosperm structure is observed by the semi-thin sections with scanning electron microscopy (SEM). Physicochemical changes in the central part of T2 endosperm are detected the by electron microscopy (TEM) and iodide staining (I_2_-KI) observations in 2022.

### A series of the quality characters and brown rice phenotypes detection in N2/N3 mutant

The phenotypes of the brown rice were detected, including the shape of brown rice such as grain length, grain width, grain thickness, and the chalky phenotype of the milled rice per plant are detected such as chalky ratio, chalky aera, the rice broken ratio. All the characters are evaluated according to the standards of key lab in China, there are 3 samples randomly selected from 100 g seeds using for the phenotype detections. The chalky characters during the Ko-mutant endosperm development are also detected by the observations of semi-thin sections and hand-cut sections in floury endosperms (3 DAP/5 DAP).

### Physicochemical properties detection in milled rice

The milled grains of the different mutants were dehydration under the ordinary temperature in our lab (28°C-30°C) and the percentage of grains with chalkiness (PGWC), the area of chalky endosperm (ACE), the degree of endosperm chalkiness (DEC) were detected, Meanwhile, the physicochemical properties such as protein content, the total starch, amylase (AS) /amylopectin (ISA), the proportion of chains (DP), and the SSC including sucrose, glucose, fructose. All the physicochemical properties of milled rice mutants compared with wild type were calculated by the student t-test (*p* < 0.05).

### The relative expression in T1 seedlings by qRT-PCR

In 2021, T1 generation of all the mutants were planted in the field with 5 x 3 lines arrangement, the total RNA was isolated from the specific tissues in whole plant (95 DAG, days after germination) are collected directly in the field, such as root, stem, sheath, flag leaf and seeds (3 DAP). The total RNA of these tissues from the whole transplant are extracted by TRIzol (Invitrogen, CA, USA) for the reverse transcription to total cDNA of plant.

The house-keeping gene ubiquitin (LOC_Os03g13170, mRNA accession number AK100267) is used as an internal control in qRT-PCR, three biological replicates were performed and the primers of qRT-PCR are listed in **Table** S13.

### The total RNA extraction and library construction for RNA-seq

For the RNA-seq analysis, the total RNAs of panicles (3 DAP) were extracted by total RNA Mini-prep kit (Axygen, http://www.axygen.com), the PrimeScript Reverse Transcriptase kit (TaKaRa, http://www.takara-bio.com). And the quality of total RNA was detected by NanoDrop ND-1000 (NanoDrop, Wilmington, DE, USA). The isolation of total RNAs by using the DNA kit. According to the manufactures’ instructions including two rounds of amplification by using a (RNA extraction kit) Message Amplification II R (Thermo fisher, https://www.thermofisher.cn/).

RNA-Seq libraries were pooled and sequenced on a lllumina nextseq 500 platform (illumine, http://www.illumina.com/). All the sequencing database were performed as the raw data according to manufacturing instructions by the Illumina Nextseo 500 control software.

### Data analysis of transcriptome and proteome for co-network construction

The RNA-Seq analysis was based on seeds (3 DAP) of all the mutants T2 planted in the summer of 2021 without treatment, and two biological replicates for each mutants were used. The raw data is obtained from the illumine-seq machine. The clean data was aligned to the reference rice genome MSU.7.0 (http://rice.plantbiology.msu.edu.). According to **the Rice Genome Annotation Project**, the criteria for data filtering and clustering are at least 6 out of 10 samples which have significant P-values, so as to make sure the normalized data qualified. After deleting redundant sequences in the raw data, heatmaps of DEGs and Venn diagrams were constructed for the genes that control seed quality.

The significant DEGs were identified and screened by using the actor. The heatmap diagrams and DEGs (log_2_FC) are verified by the quantitative PCR analysis which are performed as another format of DEGs expression.

The PCA, volcano plots and Venn diagrams are based on gene expression classification, and the GO enrichment and KEGG pathway analysis are based on the expression pattens of genes that are significantly changed between mutants and WT. All these RNA analysis visualizations are running on the platform of the LC-BIO company (http://www.lc-bio.com).

In this study, the Mapman framework of Starch and Sucrose metabolic pathway was constructed, however, the results display in a new way. As all the DEGs data are based on the RNA-seq KEGG, the ECs in KEGG are used for the hierarchal allocated DEGs in Mapman functional categories. The pathway analysis was performed through Heatmap and qRT-PCR verification for each ECs in the Mapman KEGG pathway (Fig. 5B) [26]. The primers of these genes using in qRT-PCR verification are designed by PrimerPremier 6.0 (Table. S12). All the diagrams related to the basic analysis of RNA-seq are constructed by the software TBtools 11.02 [27].

The **STRING** analysis of Pro-Pro interaction are performed by “annotate your proteome to STRING” section, which can be used for the transcription factors such as OsNAC02, and its regulated co-network proteins as well. The proteins in pro-pro subgroup with DEGs enrichment (threshold_score > 900) are selected for STRING construction in mutants versus WT (STRING, https://cn.string-db.org/). Submit the proteome of DEGs, STRING will output the interaction network and predict the protein functions (including GO terms, KEGG pathways).

### The yeast-one-hybrid assay

The OsNAC02 with its effective genes in the pro-pro interactions are proved by the yeast-one-hybrid (*Saccharomyces cerevisiae*, Y-1-H) from the Wan academicians’ laboratory. This system are including *pB42AD* vector as the active part and *pLacZi* vector as the effective part. The CDS of *OsNAC02* was cloned into the *pB42AD* vector (EcoR I), and the promoters sequences (-2000bp-’ ATG’) of the relative genes are linked with the *pLacZi* vector (Xho I), including OsMYB04, OsHSFA2E and OsHSFA2C. The positive control was constructed by the reference information relative to the key genes in ABA pathway[28]. The vector transformation and screening procedures were performed according to the procession in ‘WanJianmin lab’ (accession number AK100267), which was used as an interna control [29]. The primers of all the vectors construction are designed by PrimerPremier 6.0 (Table. S1).

## Results

### K0-*OsNAC02* /*OsNAC06* mutant construction and the genotype detection

In this study, the functions of *OsNAC02* (Os02g0594800, LOC_Os02g38130) and *OsNAC06* (Os06g0267500) were mainly identified by Ko-mutant construction. However, functional detection was performed through the online bioinformatics analysis and RNA-Seq analysis processing in mutants, which provided ways to discover their functions and revealed various phenotypes in response to abiotic resistance as well as the prematurity which may seriously reduce the quality of seeds.

The multi-mutant of N2 *mut* (*Osnac02*) and N3 *mut* (*Osnac02/Osnac06*) was created by CRISPR/Cas 9. All of the mutant sites were confirmed through PCR sequencing (Table 1), and the leaves of N2/N3 *mut* (T2) were selected for the mutant detection in detail. The N2 *mut* has two mutations in the CDS region of *OsNAC02*, while the N3 *mut* has one mutant sites in the CDS regions of *OsNAC02* and *OsNAC06* respectively (S1 **Fig.**).

**Table 1.**
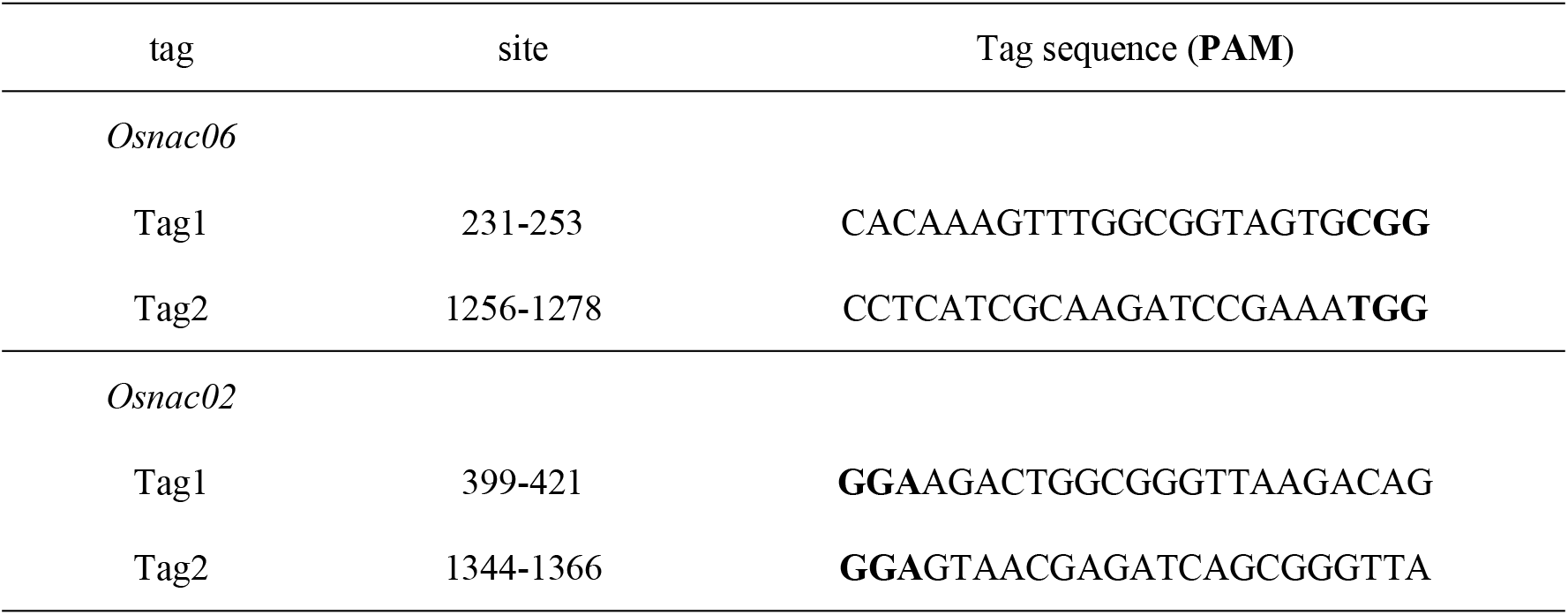
The tags of target sites designed for Osnac02 /Osnac06 mutants

Premature seed development is presented in both N2 and N3 mut (95 DAG). In the summer of 2021, N2/N3 mut seeds (T2) were harvested at 95 DAG, and the panicles matured early obviously. The period statistics of endosperm development showed that the N2/N3 mutant matured 10-15 days earlier than the WT (Fig. 1D). At 95 DAG, the caryopsis phenotype in the last stage of the grain-filling period differentiated substantially more than the WT. We found that the WT caryopsis was still green while all of the mutant caryopses were in a yellowish state.

Meanwhile, the shape phenotypes of seeds at 95 DAG and 125 DAG are calculated separately for the maturity verification. The seeds (95 DAGs) of N2/N3 *mut* have superior length and thickness of caryopsis than WT (Fig. 1E), but become incompletely filled in the mature seeds (125DAG) (**Fig.** 1F). And all the botanic morphology and yield characters of N2/N3 *mut* (T2) are detected in 2021 (S2 **Fig.**).

### *OsNAC02* (Os02g0594800) expression pattern and phylogenetic analysis of OsNAC family in rice

In this study, qRT-PCR investigated *OsNAC02* was used to investigate the relative expression of *OsNAC02* in all of the tissues of the seedlings, including the root, stem, sheath and flag leaf (45 DAG). **Figure 2A** shows that the relative expression level of *Osnac02 mut* and *Osnac06 mut* based on qRT-PCR analysis were in accordance with those on the Plant eFP Viewer (http://bar.utoronto.ca/eplant_rice/) (Fig. 2B)[30], both were overexpressed in the root, SAM and the panicles during the early stage of development.

There are a total of 195 genes containing the NAC domain in rice, and 133 genes were selected for NAM phylogenetic analysis, and these are as same as the analysis in 2003[16] (S3 Fig.) there are 133 OsNAC genes are clustered into five subgroups. *OsNAC02* (Os02g0594800) belongs to the subgroup III and *OsNAC06* (Os06g0267500) is clustered into subgroup IV having only seven genes. The homologous genetics of *OsNAC02* and *OsNAC06* were highly conserved (S3, S4 Fig.), and these genes were conserved in almost all of the plants, then *OsNAC02* were identified through homologous analysis with orthologous genes from rice, maize and Arabidopsis. *OsNAC02* (LOC_Os02g38130) and *OsNAC06* (Os06g0267500) were aligned with other seven proteins: Maize-NAC, NAC1, NAC2, ANAC044, ARATH-NAC085, NAC_044 and At1g26390, and the clustering results indicated that the N-terminus of the protein domains (0-244 aa) was highly conserved, but the sequences changed variously on the C-terminus (S4A Fig.). The location of the motifs of the promoters in the relative genes are primarily homologous in rice. The *OsNAC02* regulated genes including *OsHSFA2C*, *OsHSFA2E* and especially OsMYB04 in the PAL pathway, are highly conserved in all plants (S5 Fig.).

*OsHSFA2C* and *OsHSFA2E*-related heat shock proteins were found to be directly regulated by *OsNAC02*, as well as *OsMYB04*. These downstream transcription factors are multiple effector responses to both abiotic stress and other hormonal regulation. All of these genes containing “ACGT” motif, such as the G-boxes and the ABRE motifs in the promoter region, and their transcriptional functions were determined by the multiple effective motifs and regulatory genes. Among the many cis-elements, the ABRE/G-box motifs located in the promoter of *OsNAC02* effective genes were collected and listed (S5A Fig.).

Meanwhile, the regulation of OsNAC02 with relative TF genes can be dedicated by comparison of qRT-PCR results in root and leave of seedlings in mutants (45 DAG). As **Figure** 2D shows, the expression of all the TFs in low-level was consistent with OsNAC02 losing function in root, but normally expressed in leaves which is accordance with OsNAC expressing lower with lower affection in root. As **Figure** 2C shows, all the genes are positively regulated by the OsNAC02 directly, and their expression level keep the same trend of the OsNAC02 expression in all parts of tissues. But the expression of *OsHSB2c* in root is higher than other genes, the reason especially for which may be regulated by OsNAC02 directly.

### Chalky phenotype and quality characters of brown rice in mutants

The grain chalkiness was detected in three repeated samples selected randomly including the percentage of chalky grain, percentage of are chalky area, besides the rice broken ratio (S6A Fig.). The rice chalky ratio of N2 *mut* was mostly significantly high, and the yield of N2 *mut* was lower than that of the WT (S2C Fig.). Moreover, the yield of N3 *mut* was higher than that of WT, while the rice broken ratio of the N3 *mut* double mutants was most significantly among all materials (S6A Fig.).

As shown in the Figure S6B, the milled rice of N2 *mut* has floury endosperm a with white-core and white-belly appearance, and the chalky phenotype was more obvious than N3 *mut* (Fig. 3A). Moreover, the chalky phenotype of the N3 *mut* milled rice was similar to that of WT in 2021. Both N3 *mut* and WT showed a small white-core chalky phenotype at the center of the endosperm which was caused by the persistent high temperature in summer of 2021, but the decrease in chalky rate may not rescue the quality of N3 *mut*, for the significantly fragile phenotype with more than an 85% broken ratio in N3 *mut* milled rice (S6C Fig.).

**Fig 3.**
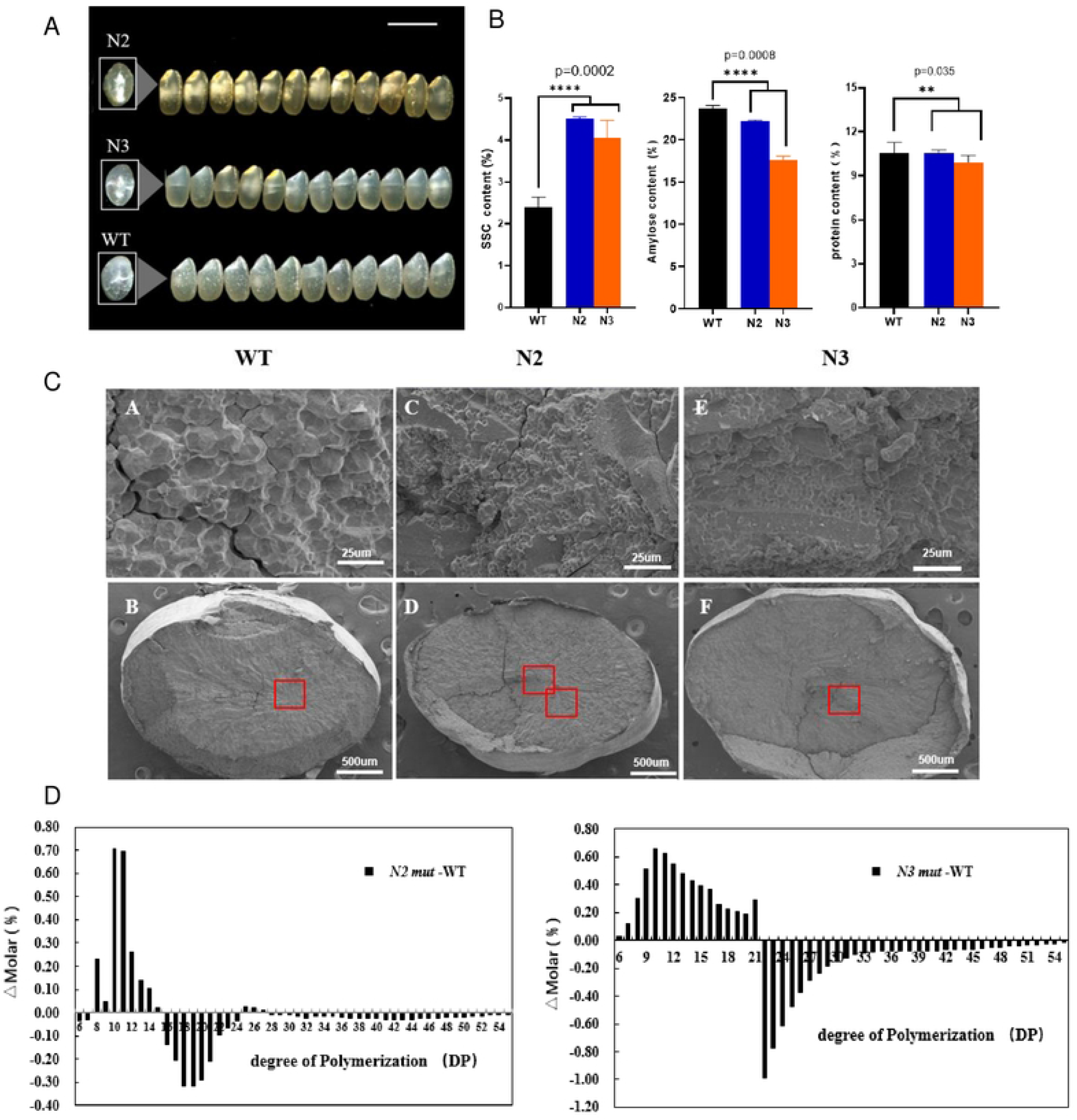
The chalky phenotype and physiochemical analysis in the N2/N3 mutants vs WT. All the characters in this figure are detected in full mature seeds (125DAGs). **(A)** The chalky phenotypes and shape phenotypes of N2/N3 mut compared with WT. All the T2 mutants are planted in Fuyang (Hangzhou, 2021), n=15mm. There are obvious white-core and white-belly in N2 mutant which chalky area is the largest, while there is medium transparent in WT and white-core only in N3 mutant. **(B)** The physiochemicalanalysis of N2/N3 mutants is distinct compared with WT. The quality characters such as SSC, amylose content, proteincontent were tested and the results were as follows: The SSC was highest in N2 mutant but lowest in WT; The amylose content was lower in N2/N3 mutants compared with WT, which was reversed to the SSC tendency in mutants; The protein content was lowest in N3 but no significant difference between WT and N2 (two-way ANOVA for mixed model, *, p < 0.05 and **, P < 0.01). **(C)** The hand-cut sections of full-mature endosperms are scanning by SEM (scale bar, 25 mm and 500 um). The starch granules arranged distinctly between N2 and N3 mut compared with WT, and the structure of starch granules are smaller and more polyhedral in mutants. (D) The amylopectin analysis shows the apparent difference in N2/N3 mut and WT. The chain lengthdistribution of amylopectin analysis was calculated by the number of short chains with degree of polymerization (DP).There are more short fragments in N2 mut while more medium fragments in N3 mut.

Overall, the chalky area and ratio of N2 mut were significantly higher, the rice broken rate of N3 mut was mostly increased, and the phenotypes analysis of mature seed (125 DAG) such as length, width, and thickness of seed was in the tendency of WT > N2 > N3 (Fig. 1F). Moreover, the yield per plant of T1 mutants (125 DAG) was in the order of N3 > WT > N2 in 2020 (S2C Fig.), but N2 > N3 > WT at the 95 DAG of T2 mutant in 2021 (S2E and S2F Fig.).

The hand-cut sections of the matured endosperm (125 DAG) were scanned by scanning electron microscope (SEM) (Fig. 3), the results showed that the starch granules were arranged distinctly in the mutants compared to that in WT. The number of starch granules was greater in N2/N3 *mut*, which developed into a floury endosperm. The volume of starch granules was smaller in N2 *mut*, and several narrow fissures were observed among the parenchyma cell. However, the N3 *mut* granules have a faster filling rate than that in N2 *mut*, leading to a tighter arrangement of the cell wall with nearly no crack in the endosperm. However, the starch granules were round, smooth which is different from that in WT (Fig. 3C). As in the normal control, the parenchyma cells of the WT were almost flat, and the starch granules were compact with every cell wall having sharp edges (Fig. 4C).

**Fig 4.**
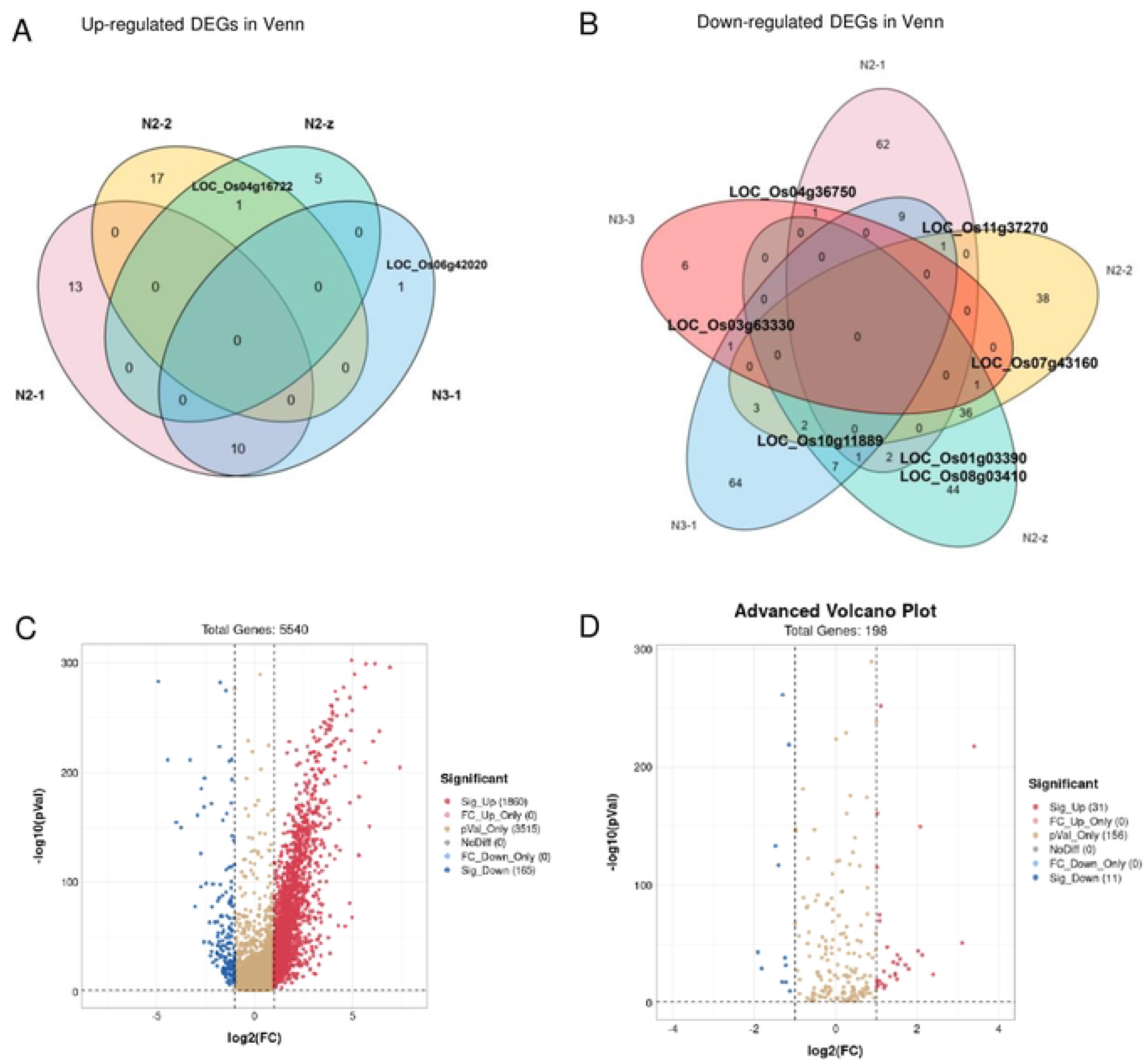
The DEGs clusters and GO enrichmentof RNA-Seq in mutant vs WT seeds (3 DAP) **(A)** The Venn diagram of up-regulated OEGs are basedon the RNA-Seq of seeds (3 OAP). **(B)** The Venn diagram of down-regulated OEGs based on the RNA-Seq of seeds (3 OAP). All the genes with label are listed in **Table S5**. In total, five mutants N2-1,N2-2,N2-z, N3-1,N3·3 were used for the cluster analysis, and the N2-2 and N3-1 duplicated samples are N2-z and N3-3. **(C)** The volcano cluster of the OEGs in RNA-Seq of seeds (3 OAPs). As cluster shown, most of the OEGs in red are up-regulated genes, the down-regulated genes are in blue and the genes expression with no difference are in yellow. All the genes expressed significantly are listed in **Table S8**. **(D)** The volcano cluster of the OEGs in Sucrose and Starch pathway of RNA-Seq in N2/N3 mut vs WT seeds (30APs). All significantly expressed genes are listed in **Table S10**.

The iodine-stained semi-thin sections of the opaque endosperm in LM (5 DAP, light microscope) showed that amylose content (AC) was higher in N3 *mut* than in WT versus N2 *mut*, which could be verified examining the darker blue color in the early endosperm. Furthermore, as shown in Figure S7 for the opaque endosperm scanned by the TEM, N3 *mut* showed more compact intercellular locking of parenchyma cells than N2 *mut*, and the parenchyma cell volume was larger and even overextend the space between the cells (S7L Fig.).

### Physiochemical analysis of milled rice in mutant vs WT for quality characters verification

The chalky phenotype of the mutant seeds was detected after ripening. In 2021 summer, all of the materials were planted in the field (Fuyang), and there was a higher temperature during the grain-filling stage. The brown rice had chalky phenotypes, such as white-core and white-belly in the N2 *mut* endosperm obviously. In addition, the fragile phenotype of milled seed in N3 *mut* was obvious. For the significant distinction with of the WT, a serious of indexes for physiochemical analysis were detected in our lab, including: the soluble sugar content (SSC), amylose content (AC), protein content (PC), and amylopectin chain length distribution analysis – the degree of polymerization (DP) (Fig. 3).

The SSC was significantly higher in N2/N3 *mut* than in the WT, while the AC decreased significantly in N2/N3 *mut*; however, the soluble protein content hardly changed (Fig. 3B). As shown in Figure 3D, according to the length distribution of amylopectin analysis, the DP in N2 *mut* showed that the number of 8-14 small fragments (DP) was significantly greater in the mutant than that in the WT, while the number of ultra-short chains with 6-7 (DP) and the number of medium and long chains with 16-24 (DP) significantly decreased during the same time. The number of short chains with 6-21 DP in N3 *mut* was significantly higher than that in the control, while the number of long chains with ≥ 22 DP was significantly lower than that in the control. This trend is different from that in the N2 single mutant (N2-1 *mut*). As shown in all of the results below (Fig. 3D), according to the different chain length distributions of endosperms, there were obvious differences in the chalky phenotypes and starch particle size (Fig. 3C).

### PCA of RNA-Seq and heatmap in mutants versus WT seed (3 DAP) in 2021

As the Principal component analysis (PCA) of seeds (3 DAP) of N2/N3 *mut* in 2021 shown that the WT and N2-1 *mut*, N2-2 *mut*, and N3 *mut* were distributed significantly, and they formed three clusters independently in the 2D-image of the PCA (S8 Fig.). This means all were applicable and qualified for the RNA-Seq analysis; the DEGs were more credible in this study.

Based on the results of the RNA-Seq analysis in **Figure** 5, the heatmap analysis of the N2 mut seeds showed that the expressed level of the DEGs were lower than those of DEGs in the WT, while the expressed level in the mutant distinct genes in N3 mut were higher than the DEGs in the WT, and the expression level of most genes were in the order of N3 > WT > N2.

**Fig 5.**
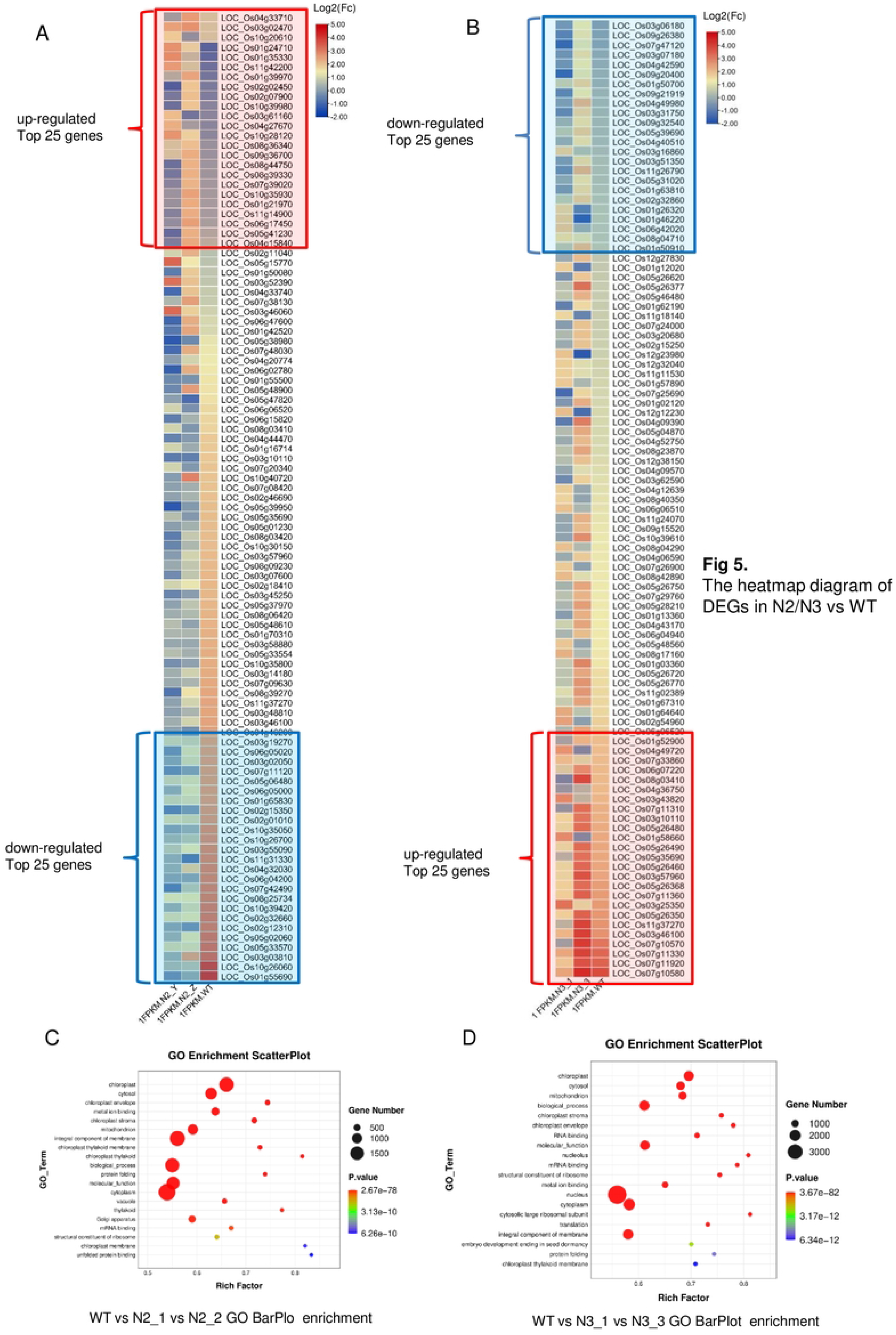
The heatmap diagram of DEGs in N2/N3 vs WT. **(A)** The distinct expressed genes (OEGs) in the heatmap of **N2 vs WT** in an ascending ordering; **(B)** The distinct expressed genes (OEGs) in the heatmap of **N3 vs WT** in an ascending ordering, all the RNA-seq data were standardized by the log10 parameter, and the top 25 genes with highest expression are highlighted in the red frame, and the top 25 genes with lowest expression are outstanding in the blue frame; **(C)** The GO analysis of the **N2 vs WT** was according to the top 20 GO enrichment pathways of **N2-1 vs N2-2 vs WT**; **(D)** The GO analysis of the **N3 vs WT** was according to the top 20 GO enrichment pathways of **N3-1 vs N3-2 vs WT**

The top 25 genes with low expression-levels in the mutants (blue region) seeds are mainly related to pathogen invasion resistance, protein in the root tip for heavy metal ion resistance, and salt stress response. Four genes are related to disease resistance, including laccase synthesis precursor (lac15), cyclosporin and plant lectin. Previous studies indicated that the PAL pathway was one of the most crucial synthesis pathways for tea (*Camellia sinensis* L.) to respond to gray mold disease stress, and LAC15 (TT10) was the primary enzyme for anthocyanin synthesis [31, 32]. The expression of the mutant was higher than that of WT, indicating that anthocyanin synthesis was upregulated in the PAL pathway. The key enzymes of the PAL pathway in turn include: CESA, CHS, CYP, ERF96, ETI, FLS, lac15, PPO, PR (the pathogenesis-related protein), TLP and UDPG. The OsI-BAK1 protein contains the eLRR-domain for the resistance of the brown Planthopper Nilaparvata lugens [33].

On the contrary, the expression level of top 25 genes (red region)was higher in N2 mutant than that in WT, as follows: the seed gluten synthases (2×), enzymes (5×) related to the starch synthesis pathway including the sucrose synthase, starch synthase, glucan phosphate acyltransferase, 1,4-glucan branching enzyme and pyruvate; the TFs, such as *dof* Zinc finger, *OsNAC023* and *OsNAC025*; the intracellular mitochondrial transporters related to energy conversion (1×), pyruvate kinase (1×), AGPL (1×), PPI phosphorylase (1×), and early Nodulin forming proteins (2×); abiotic stress-related proteins such as metal-ion transporter (1×), channel protein (1×), CAMK (1×), the stress response protein (1×), and the Nudix hydrolase family (1×) (Fig. 5A and Table. 2).

The heatmap analysis of the N3 versus WT seeds showed the expression level of top 25 genes upregulated in the N3 *mut* versus WT (Fig. 5B, red region), these nine genes are including seed Gliadin synthesis (9×), seed gluten synthesis (1×), seed storage protein inhibitor (ITPL, 4×), two genes related to pectin synthesis inhibition (2×), heat shock protein chaperone (HSP 20, 1×), mycoprotein (AMBP1,1×), and the Cupin family proteins (3×). Contrary, the top 25 genes in the WT expressed at low level (blue region) were expressed lower in N3 *mut* compared with WT, as follows: genes related to starch synthesis including β-amylase, late embryogenesis abundant, late embryogenesis abundant protein; glycosyl hydrolase (1×), starch binding protein (GBP) and polymerized synthetase 3; dehydrated protein synthetase (2×), superoxide reductase (Aldehyde ketone reductase, 1×), endosperm development forming protein (3×), ion transfer protein (1×) (Table. S3). The functional protein *GD1*(LOC_Os09g32540) related phenotype formation was downregulated in the DEGs of N3 versus WT. LGD1 has one copy in rice; it is unique to *Gramineae* only and mainly located in the nucleus, and it regulates the vegetative growth and development of rice through its specific spatiotemporal expression-pattern and protein-RNA binding activity of multiple transcripts [34]. In previous research, the heat shock protein chaperone HSF7.1 (LOC_Os03g16860) was expressed stably under heat stress, while all the other genes were expressed at low-level. In this study, HSF7.1 expression increased in seeds of N3 *mut* under normal conditions, which was not in accordance with the results of a previous study results and it may indicate that the deletion of *OsNAC02 /OsNAC06* will remove the inhibited factor from the HSF7.1 [35, 36].

Based on the RNA-Seq seed results, the significant DEGs of mutant and the WT were selected, and 75% of them were related to the starch metabolism, and the rest were related to the monosaccharide and energy metabolism pathways. Overall, the significantly expressed DEGs in the seeds of RNA-Seq was mainly related to the starch metabolism pathway, and 90% of these genes were up-regulated, which were related to the starch metabolism pathway in the leaves and roots of mutant seedlings. The expression levels of genes in the N2/N3 mutant were at low-level, and they were even lower than those in the WT. These genes include the terpene biosynthesis enzyme, glycosyl reductase, and Anth/Enth protein, which are mainly involved in endocytosis mediated by the AP2-independent Clathrin protein. All of the functional proteins involved in biological disease resistance were downregulated. Meanwhile, the proteins related to plant hormones such as regulation auxin induced proteins and BRI receptors were downregulated (Table. S2).

### Gene ontology (GO) and KEGG pathway enrichments in the seeds

As shown in the Gene ontology (GO) analysis of RNA-seq in **Figure** 5 and S9, among all of the GO enrichment pathways in the N2 versus WT seeds (3 DAP, 2021), most of the DEGs in the KEGG pathways were significantly enriched in the chloroplast and chloroplast thylakoid, which is in accordance with the results in **Figure** S9C. Many of the genes enriched in the GO pathways had a lower rich factor (R < 0.06), including the cytoplasm, chloroplast envelope, integral component of membrane, biological_process and molecular function. In **Figure** 5D, the GO enrichment pathways of the N3 versus WT seeds, showed that more distinctly expressed genes in the GO pathways were significantly enriched in the chloroplast and mitochondrion compared to those in N2 versus WT, which is in accordance with the results in **Figure** S10E. Meanwhile in **Figure** 5C and 5D, many other genes with less rich factors (R < 0.06), which were still enriched in the cytoplasm and nucleus pathways.

In **Figure** S9C, the GO bubble statistic of the biological process (blue bubble), cellular component (red bubble), and molecular functions (green bubble) are in accordance with the results in **Figure S9A** and **S9E**. Thus, all of the GO enrichments in **Figure S9C** for N2 versus WT, the GO bubble (GO:0009507, chloroplast) and (GO:0009535, chloroplast thylakoid membrane) were in accordance with **Figure S9A** with the green part of GO results: there were 1749 genes in GO :0009507 and 236 genes in GO:0009535. While in **Figure S9D** for N3 versus WT, the GO part: red bubble (GO:0005739, mitochondrion) and green bubble (GO:0005618, cell wall), which matched the results in **Figure S9B** with the GO there were 1171 genes in GO:0005739 and 218 genes in GO: 0005618 (Table. S4). **Figure S9E** represents the top 20 GO pathways of the N2 *mut* versus WT seed transcriptome, including three cellular component pathways: GO:0009507 is placed in the top 1 as the chloroplast pathway, GO: 0009535 is placed in the top 20 as the chloroplast thylakoid membrane pathway, and GO:0005739 is placed in the top 3 as the mitochondrion pathway. As confirmed in the GO enrichments, N2 *mut* displayed a distinct chalky phenotype, mainly due to significant enrichment of the related expressed genes in the chloroplasts, chloroplast thylakoid and mitochondrion, which control energy metabolism (S9C, S9E Fig.).

Energy metabolism is the conversion from ATP to ADP, Pi. Extracellular ATP (eATP) is involved in the GO:0009535 pathway. In previous study, the nanomolar levels of eATP were accumulated by wounding in the plant, which triggered the ROS wave during the wounding process [37]. Meanwhile, eATP is secreted from extruding cells during the pathological invasion or epithelial homeostasis disorders, which promote the ROS level for the biotic resistant [38]. The cell walls of the N3 *mut* seeds are remarkably fragile, thus resulting in a very high rice broken rate, which is mainly related to the cell wall pathway (GO: 0005618). Meanwhile, the genes related to starch synthesis were down-regulated in N2 *mut*, which changed the synthesis pathway and function (Fig. 7C). Moreover, these significantly enriched genes in GO are related to cell wall synthesis such as *OsCIN*, which plays a role in inducing defense in rice through sugar homeostasis regulation[39, 40]; There were no qualitative changes in the storage of the endosperm, which implied that N3 *mut* may have a recovered mutant loci for the qualitative character but N2 *mut* does not. Meanwhile, the formation of cell walls underwent changes astonishingly, resulting in the rupture of individual cell walls under slight external mechanical pressure in N3 *mut*, which may have changed the structure of the endosperm and led to fragile phenotypes (Fig. 3).

Although the phenotype of the broken rice is commonly found in rice harvests, there are still insufficient theoretical studies on these reasons. According to the heatmap of N3 versus WT, we revealed that the DEG expression in N3 mut was significantly lower than that in WT (Fig. 5B, blue region), mainly including: ZmEBE-1 (LOC_Os01g26320), GDSL-like synthase (LOC_Os01g46220) and CSLA9 (cellulose synthase 9, LOC_Os06g42020) in cytosols. It was confirmed that the three genes involved in endosperm formation in N3 mut, are related to cell wall synthesis and endosperm filling (Table. S3 and S5). The expression of all the genes involved in cell wall synthesis was downregulated, causing the fragile phenotype with significantly higher broken ratio in N3 mut.

### Venn diagram in N2/N3 *mut* versus WT seeds and the volcano plots of DEGs in starch metabolism in the N2 *mut* versus WT seeds

The Venn diagram shows that the DEGs were in upregulated and downregulated groups, which were based on the top 99 DEGs in the seeds (3 DAP). As shown in **Figure 4A** of the Venn diagram, 90% of upregulated genes in N2-1 versus N3-1 groups were co-expressed genes, except for the *CSLA9* (LOC_Os06g42020), and upregulated DEGs in each mutant were clustered individually. This result means that all the upregulated DEGs of the mutants have been enriched in one metabolism pathway that was most active during early endosperm development.

Comparing all the down-regulated DEGs in the overlaps of Venn diagram in mutants (**Fig. 4B**), we found that 50% of the genes in N2-2 mutants could co-express with other mutants, while other mutant DEGs were clustered individually. Through these co-expressed genes, the main pathway which was unique in the mutants could be discovered. There are two genes in the overlap of N2-1 versus N2-2 mutants, *BBTI7* (LOC_Os01g033900) and *glutelin* (LOC_Os08g03410) were co-expressed and involved in the quality formation processes such as the regulation of energy storage and the accumulation of endogenous seed proteinases as well as thiamine. At the same time, both genes are involved in defense systems that mainly respond to biotic stresses, such as pathogens, insect bites, and mechanical damage, and they are highly expressed in flowering and early endosperm formation[41].

Through the downregulated DEGs, aspartokinase (LOC_Os03g63330) was co-expressed in both the N3-1 and N3-3 mutant, which was involved in the amino acid synthesis in response to drought stress [42]. The protein partner HSP20 (LOC_Os04g36750) co-expressed in N2-1 and N3-3 mutants, was mainly involved in the activation process of the stress-induced abiotic stress response [43]. All of the above genes mentioned are co-expressed in the N2/N3 mutants. Overall, there were fewer co-expressed genes in the downregulated DEGs of Venn diagram in N2 and N3 mutants. The co-expressed genes in all of the mutants were all related to biological stress response (Table. S4).

One conserved protein (LOC_ Os10g11889) is a functional gene related to plant biotic stress response, and it is co-expressed in three N2 mutants (**Fig. 4B**). Although this protein domain is highly conserved, a series of these genes, which functions under rice blast and pest stress conditions have not been reported yet [44]. *AMBP1* (LOC_Os11g37270) is co-expressed in N2-1, N2-2 and N3-1 mutants (Fig. 4B), and it could act on invasive disease by synthesis of chemicals that have a broad-spectrum and high-efficiency resistance. According to the co-expression in tissues throughout the plant, this gene is specifically expressed at a higher level on the embryo, including the vegetative embryo and early endosperm post-anthesis. However, *AMBP1* remains an unknown gene for us as its range of functions has not been reported yet. In the rice population inferred to study abiotic stress resistance, it was only found at the candidate region for salt and alkali resistance QTLs[45].

*OsTPP3* (LOC_ Os07g43160) was co-expressed in N2-2, N2-z and N3-3 mutant, which responds to abiotic stress by catalyzing with trehalose as substrate. According to tissue-specific expression analysis, OsTPP3 was co-expressed in various tissues, but its expression level is significantly higher in vegetative flowers and seeds. Meanwhile, this gene was significantly expressed in the roots and buds of seedlings (7 DAG) under drought and salt stress treatment, and its expression level was upregulated significantly during growth and development period under stress conditions [46]. OsTPP3 has three conserved domains to form catalytic sites for this enzyme. The motif elements in the promoter are include ABRE (4×) and Motif IIb (1×) elements, which are involved in ABA response, and the HSE (1×) and G-box (7×) motifs are involved in the heat stress and light responsiveness.

In conclusion, most of these genes co-expressed in the mutants are involved in stimulus processes of the resistance response to harsh environmental stress. The trade-off balance in the energy cycles obviously affected the synthesis of the storage components of the vegetable endosperms, such as starch granules, proteinase, globulin and prolamin, which led to white-core or white-belly in the seed filling process.

Almost all of DEGs in the volcano plot of the N2, N3 versus WT were upregulated (**Fig. 4C**). The top 20 significantly expressed genes up-regulated were labeled (S12 Fig.), the 16 genes of the 20 top genes labeled were related to starch synthesis, the functions of the top 5 genes significantly up-regulated in mutants are mainly concentrated on the photosynthesis, nucleus replication and energy production and transportation (Table. S7). At the same time, the top three downregulated genes are *OsNAC020* (LOC_Os01g01470), *glutelin* (LOC_Os02g15070) and *hsp20* (LOC_Os11g13980), are mainly related with the qualitative formations such as starch and storage protein synthesis. These downregulated genes are relative to starch metabolism, decreasing protein storage and enhancing the iron bioactivity through the membrane channels. As a result, the morphology of the grain chalkiness affects quality through the presence of opaque spots in the endosperm, which induces white-core formation as well.

The DEGs upregulated in the Sucrose and Starch metabolic pathways in seeds (3 DAP) are related to the synthesis of soluble micro-molecular. *OsBAM2* (LOC_Os10g32810), *OsSSIIIb* (LOC_Os04g53310) are the essential enzymes in the primary steps of starch degradation and synthesis, while *LTPL18-protease* (LOC_Os01g12020) was involved in lipid synthesis and pathogen invasion resistance. Surprisingly, *GBSSII* (LOC_Os07g22930) is involved in the temporary starch granules transport to sink organ and *OsGLN2* (LOC_Os01g71670) involved in the 1,3-β-D-glycosidic linkages hydrophobic action of disaccharides in the cell wall, but both genes are not expressed specifically in the endosperm, but rather in the lemmas during flowering or in the germinating seeds [47]. This means that all of the genes that were significantly upregulated are related to energy transportation, not storage for abiotic resistance and the pathogen infection resistance (Table. **S9**). As previous reports, the GBSSI and GBSSII were downregulated for the promotion of starch accumulation in the leaves of seedlings under the salt stress [48]. GBSSII functions as a mediating binding protein during starch granule development, which expressed specially in leaves for the transient starch in starch metabolism [49]. The top five genes significantly downregulated were *α-amylase* (LOC_Os06g49970), *GH1* (*Os3BGlu6*, LOC_Os03g11420), *GH5* (glycosyl hydrolase family 5, LOC_Os04g40510), *β-amylase* (LOC_Os03g22790), and *CPuORF23*(LOC_Os10g40550) in the DEGs of the starch synthesis pathway. *GH1*(LOC_Os03g11420) and *GH5*(LOC_Os04g40510) belong to glycosyl hydrolase, which is involved in the chemical defense, alkaloid metabolism, hydrolysis of cell wall-derived oligosaccharides, phytohormone regulation, and lignification. *CPuORF23*(LOC_Os10g40550) is involved in Trehalose synthesis which is related to drought resistance, *α-amylase* (LOC_Os06g49970) and *β-amylase* (LOC_Os03g22790) are involved in the diurnal rhythm of altering the primary carbon flux into soluble sugar for the abiotic resistance in leaves (Table. **S10**).

### Metabolic analysis of the Sucrose and Starch pathway in the mutant seeds (3 DAP) by Mapman framework construction

In **Figure** 6, the 29 **ECs** were labeled with alphabetic numbers in Starch and Sucrose metabolic pathway (ko00500). A total of 39 genes with 29 **ECs** were selected for the Mapman analysis sucrose pathway were downregulated (Fig. 6A). The heatmap diagram showed the qRT-PCR of 39 DEGs in Mapman KEGG in N2 mutant versus WT (Fig. 6B), The different chalky phenotypes in the N2 /N3 mutant were determined by the DEGs in the starch anabolism and metabolism pathways. *RSUS1* (LOC_Os03g28330), *1,4-α-GBT2* (LOC_Os02g32660), *4-α-DPE2* (LOC_Os07g46790), *1,3- glucosidase 12*(LOC_Os02g53200), *endoglucosidase 7* (LOC_Os02g50490), and *OspPGM* (phosphoglucomutase 2, LOC_Os03g50480) were significantly upregulated in the N2 /N3 mutant. Contrary, the *β-SnRK1* (LOC_Os05g41220), *SBE I* (LOC_Os06g51084) and *GBSS1b* (LOC_Os07g22930) were significantly downregulated in the mutants compared with the WT (S11 Fig.). Particularly in the N2 seeds (3 DAP), the AGPase large subunit 2/AGPase small subunit 1, BAM2 and ISA2 as the key enzymes in starch anabolism were expressed at low levels which may be the main reason for white-core formation and small round starch granules in endosperm of N2 *mut*.

**Fig 6.**
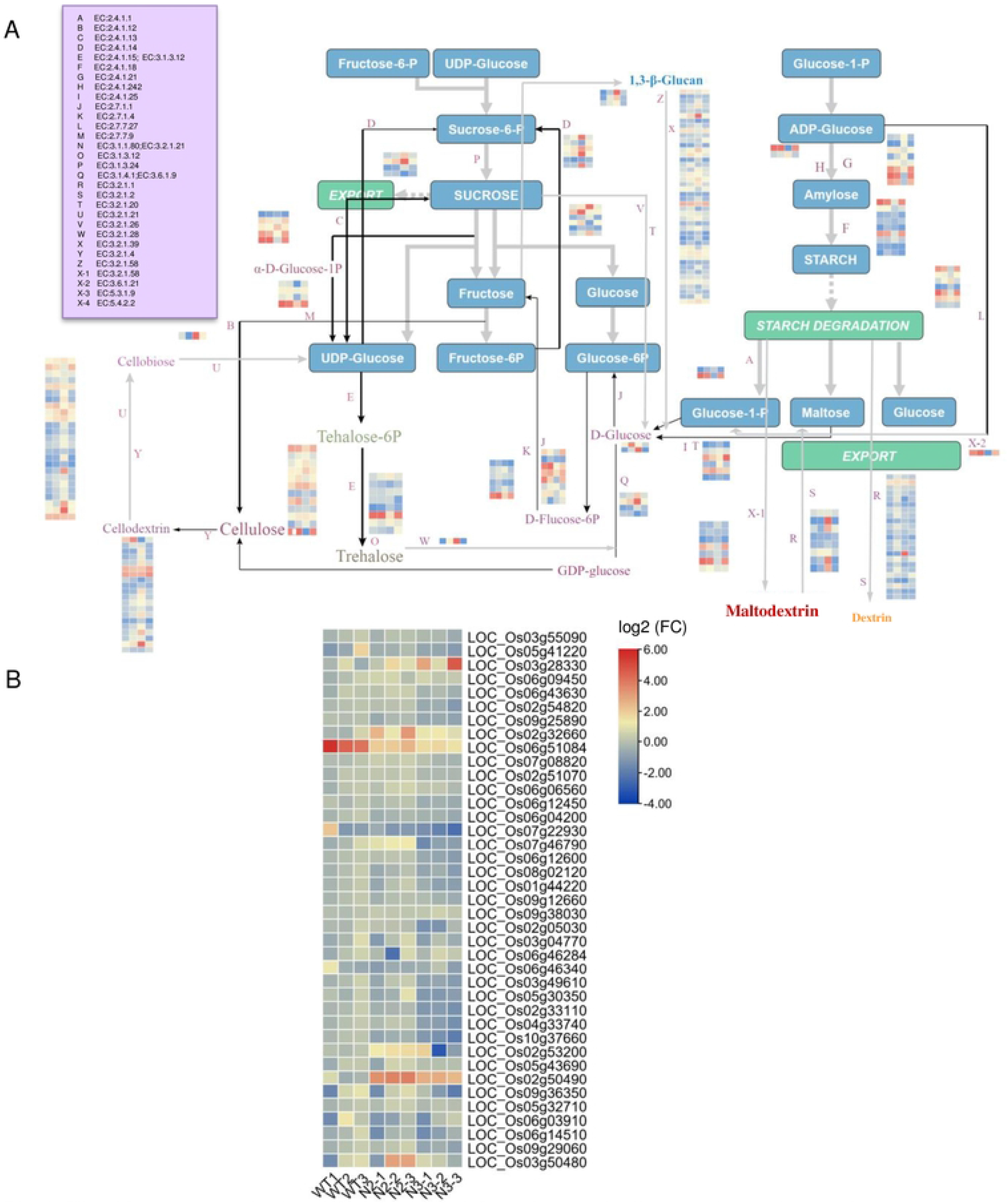
The Mapman analysis of Sucrose and Starch metabolism pathway in seeds (3DAP) of N2 vs WT and qRT-PCR verification of the DEGs in RNA-Seq KEGG. **(A)** The Mapman analyses are totally basedon the Sucrose and Starch metabolism. The DEGs are screened from the RNA-Seq of N2 vs WT seeds (3DAPs). There are 29 ECs in this KEGG pathway, and the heatmap diagram in each EC contain four samples: WT, n2-1, n3-1,n3-2 (Figure S8).This Mapman constructure shows a dynamic process related to source-sink balance duringgrain filling procession. **(B)** The qRT-PCR verifications of 39 genes significantly expressed in ECs of this MapMan, which is based on the RNA-Seq KEGG in N2 vs WT seeds (3DAPs). These 39 genes expressions in seeds(3DAP) are detected repeatedly by qRT-PCR in both N2 and N3 mut vs WT. The almost all DEGs in mutants are down-regulated while the RSUS1 (LOC_Os03g28330), OsBE2b (LOC_Os02g32660), endoglucanase 7 (LOC_Os02g50490), OsDPE2 (LOC_Os07g46790), phosphoglucomutase 2 (LOC_Os03g50480) are up-regulated in N2 mut significantly (Table S12).

Gene expression was significantly distinct in N2 versus WT: The expressions of *1,4- α-GBT2* (LOC_Os02g32660), 1,3-glucosidase 12 (LOC_Os02g53200), endoglucosidase 7 LOC_Os02g50490) and *OspPGM* (LOC_Os03g50480) were higher in N2 mutant (Fig. 6B). Most interestingly, the expression of *RSUS1* (LOC_Os03g28330) in N3 mutant was positively higher than that in N2 mutant.

### Pro-pro String screened co-network proteins in OsNAC02 *mut* versus WT seeds (3 DAP)

Based on the DEGs in the RNA-seq of N2 versus WT, there were 199 proteins of DEGs involved in the co-network construction, and only 47 proteins could interact with other proteins in forming a co-expression network (Fig. 7A). Of these 47 proteins, the most effective co-expressed proteins in the network were selected by the threshold (score ≥ 900), and three groups of significantly co-expressed proteins in the co-expression network were screened out in the N2 mutants versus WT (Fig. 7B), as follows: GIF1 (CIN2)-GIF2 (AGPL2)-Waxy-SBEI-ISA2, Tubb1-Tubb4-Tubb8 and MCM4-MCM5-RPA1B are significantly co-expressed as three groups (Fig. 7B).What we found specially, the MCM4 and MCM5 co-network proteins were significantly enriched in the N2 and N3 mutants.

**Fig 7.**
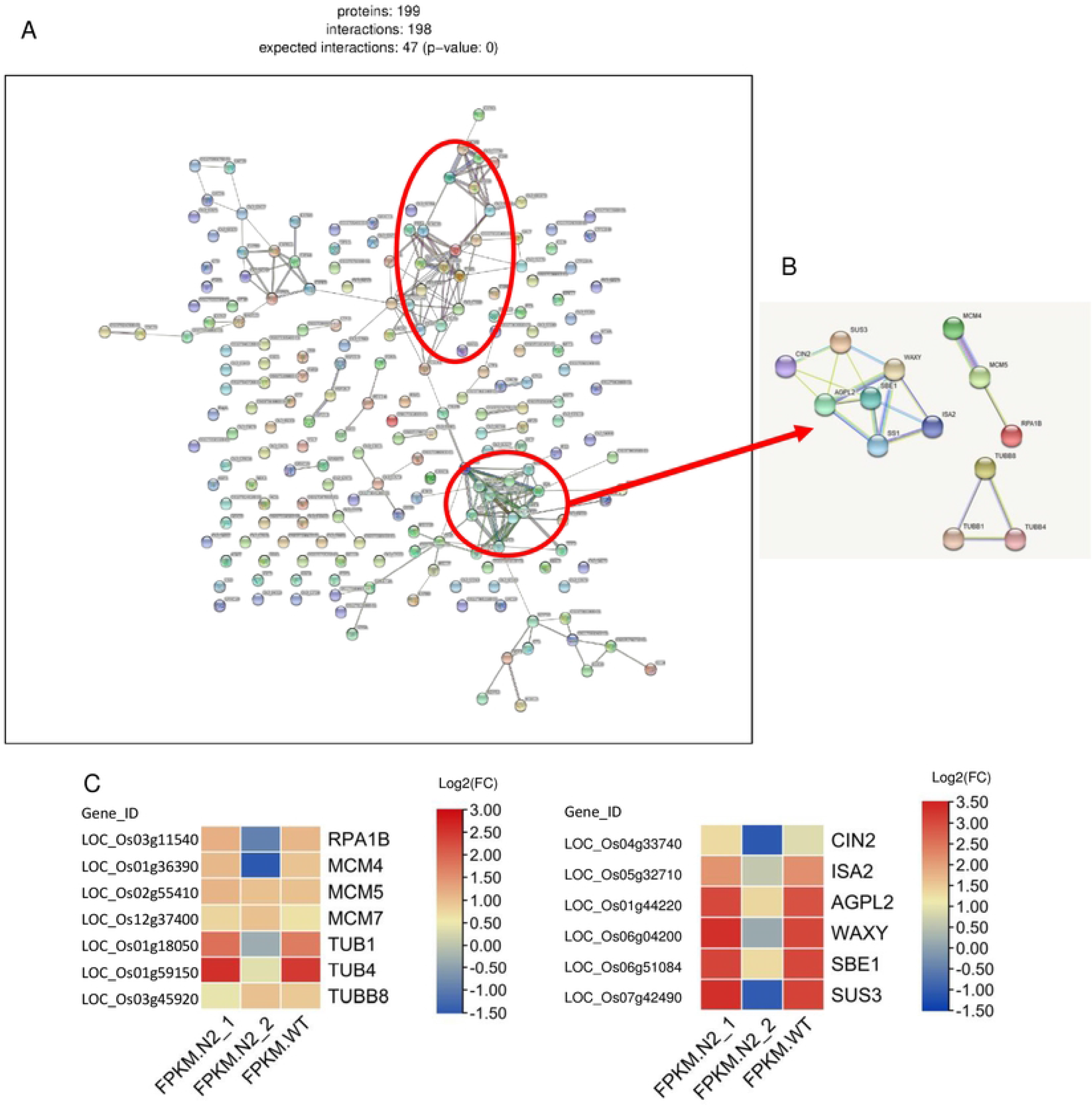
The proteins co-expressed analysis for DEGs co-network in WT vs N2 seeds(3DAPs) **(A)** In the RNA-seq of N2 vs WT seeds, all the pro-pro interactions representing the DEGs expressed in three co-networks. There are 199 proteins of DEGs involved in the co-network construction and only 47 proteins of them involved in the interaction which construct their co-expression network. **(B)** The 3 groups of proteins in co-network were screened by the pro-pro String on the Score_threshold>900, which are significantly co-expressed; **(C)** The heatmap (lg FPKM) are in accordance with DEGs expression in 3 groups of proteins which significantly co-expressed in seeds **N2 vs WT**, but SSI (LOC_Os06g06560) is not listed in the DEGs of the RNA-seq KEGG (**Table S11**).

All of these proteins could be found in the DEGs of KEGG pathway in the RNA-seq results (Table. S11), however, *SSI* (LOC_Os06g06560) was not in the DEGs of the KEGG enrichment and RNA-seq, which may be an important EC in the starch metabolism pathway of the Ko-*Osnac02* mutant, and it may be continuously expressed in a stable condition.

Eating and cooking qualities (ECQs) are determined by various reasons and this complex characteristic was controlled by a great number of genes in the co-network [50]. In this study, all of the seeds and materials were developed in the field of 2021 without treatment.

In RNA-seq results of the N2 mutant seeds, the effective and marvelous DEGs and proteins in the co-network were confirmed in the early endosperm development (Table. S11). In total, there are seven genes relative to sucrose and starch metabolism in the co-expressed group **I** (Score_threshold ≥ 900), and these genes could be arranged in a classic model of starch synthesis (S12 Fig.). *AGPL2* (LOC_Os01g44220) is a key enzyme in the primary step of the starch synthesis, and it controls the ADPG synthesis directly. This gene plays a crucial role in the regulation of starch synthesis and grain filling. *Waxy* (LOC_Os06g04200) is the key gene for Amylose synthesis, and heatmap based on the RNA-Seq KEGG of the DEGs (Fig. 7C) shows that the *Waxy* expression in N2 *mut* is lower compared to WT, whereas the heatmap based on the qRT-PCR verification of Mapman KEGG shows the expression of *GBSS I* (LOC_Os06g04200) and *GBSSI b* (LOC_Os07g22930) in N2 mutant was similar with that in WT (**Fig.** 6B).

Meanwhile, in the amylopectin synthesis pathway including: *SSI* (LOC_Os06g06560), *SBE I* (LOC_Os06g51084), and *ISA 2* (LOC_Os05g32710) were all downregulated. SSI was not significantly expressed in the KEGG pathway, but it was expressed stably in the main step of amylopectin synthesis. *SBE I* and *ISA2* were downregulated in the DEGs of RNA-Seq KEGG (Fig. 7C), which means the amylopectin content decreasing with decreasing *SBE I* and *ISA 2* expression, and then the balance between amylose and amylopectin is broken in the step of the SBE I transformation cycle (molecular amylopectin branch). As the negative feedback from the various substrates remains in the amylopectin synthesis pathway, *ADPG* will consequently flow to the amylose synthesis pathway.

Integrated all the results above with that in **Figure** 4D, as *OsBAM2* (LOC_Os10g32810) was significantly upregulated in the amylose degradation, more amylose will be degraded as the disaccharides for developing requirement (S10B Fig.). This could also be verified in the iodine-stained semi-thin sections of the opaque endosperm (5 DAP) in N2 and N3 *mut* (S7F, S7L Fig.), and the higher amylose content in vegetative endosperm of N3 *mut* than that in N2 *mut,* this result could be verified by the darker blue shown in N3 endosperm (5 DAP). *OsCIN2* (LOC_Os04g33740) is specifically expressed in immature seeds, and *gif1 (OsCIN2)* is chalky with a round smaller starch granule in the white grain [39]. In addition, *OsSUS3* (LOC_Os07g42490) corresponded to the high-temperature sensitive period during rice ripening, when its expression-level increases under the high-temperature condition; A previous study indicated that high-expression of *OsSUS3* (LOC_Os07g42490) enhances high-temperature tolerance in plants [51].

As the co-network proteins shown in **Figure** 7B, the *MCM5* (LOC_Os02g55410), *MCM4* (LOC_Os01g36390), and *RPA1B* (LOC_Os03g11540) were expressed at high levels in proliferating tissues of rice. The three genes relative to controlling the cell division in the co-network group **II**, affect grain shape and quality of seeds in mutants during endosperm development.

In this study, the *MCM4*, *MCM5* and *MCM7* proteins in pro-pro co-networks were significantly expressed in both N2 and N3 *mut*, which implies that these three genes may be regulated by the OsNAC06 directly.*OsNAC06* (OsSOG1) is the transcription factor that is homologous to AtSOG1(At1g26390) and regulates the cell cycle in the upstream [49]. It was found that genes regulate cell cycle including: *CYCB2.1*, *CYClazm*, *E2F2*, *MAPK*, *MCM5* (LOC_Os02g55410), *MCM4*(LOC_Os01g36390), *MCM7*(LOC_Os12g37400), *CAK1*, *CAK1A*, *CDKA1*, *CDKA2*, *CDKB*, *CYCA2.1*, *CYCA 2.3*, *CycD3*, *CDKD4*, *CDKT1*, *H1*, and *MAD2*.

As the stranded DNA binding complex subunit 1, *RPA1B* (LOC_Os03g11540) is a one replication protein, positively expressed in proliferating tissues such as the root tips and young leaves, which contain root apical meristem and marginal meristem respectively[50]. In eukaryotes, the MCM family of proteins plays an important role in chromosome replication process. The normal DNA replication in every cell cycle is accurately controlled by binding the origin recognition complex that recruits MCM proteins including MCM2 - MCM7, which are highly conserved in organisms [52, 53]. The MCM complex consists of six different subunits (MCM2 - MCM7), and it interacts to form a ring-shaped hetero-hexamer with a central channel large enough to encircle the DNA. Once the MCM complex has been loaded, origins are licensed to replicate and any site containing the MCM complex has the potential to form an active DNA replication fork [52]. In human beings, *MCM4* expression decrease about 5% of the normal level, which will cause the DNA damage response[54]. In Arabidopsis, the subunits of the MCM2-MCM7 complex were coordinately expressed during the development stage, which is abundant in proliferating and endocycling tissues. This displays the indicative role of DNA replication in plants, the *MCM5* and *MCM7* subunits remain in the nucleus during the S and G2 phases of the cell cycle, whereas *MCM4*, *MCM6*, and *MCM7* form a “core” complexes, which are more tightly associated with chromatin than the remaining subunits [53].

The three Tubulins such as *OsTUB1* (LOC_Os01g18050), *OsTUB4* (LOC_Os01g59150) and *OsTUB8* (LOC_Os03g45920) were interactive in the co-network group **III**. There are α-tubulins and β-tubulins in the Microtubules family, which are strictly conserved in protein coding regions of almost all eukaryotes. There are eight TUB isotypes with 27 predicted amino acid sequences of *TUBs* in rice [55].

Tubulin protein is reported to be expressed in the dividing tissues mainly[56], which may play roles in most of the cell division processes and cell elongation [57]. However, in a previous study, the tubulins (TUBB) are verified to be expressed tissue-preferentially, and they are positively expressed in dividing tissues, including root tips and tender leaves. However, the reduced expression of *AtTUA6* (α-tubulin) in Arabidopsis has been shown to affect root growth and morphogenesis severely [58]. *OsTUB4* (LOC_Os01g59150), which is expressed stably in all tissues, is used as the internal control for qRT-PCR in almost all plants [59], while the *OsTUB1*(LOC_Os01g18050) function has yet to be elucidated. Meanwhile, as an anther-specific β-tubulin, *OsTUB8* (LOC_Os03g45920) is also expressed at high levels in vegetative growth, such as anthers and seed set in rice[57]. Because these Tubulins are highly conserved in DNA sequences and play an essential role in cellular processes, their low-level expression may also affect the most active division process - the grain filling.

## Discussion

There are two phases in a plant growing period: the vegetative phase with germination and plant growth, the reproductive phase with flowing and spikelet fertility. During these plant growing processions, the sugar signals play an important role through the sucrose and starch metabolic pathway. The glycolysis dependent on SnRK1/TOR signal transduction pathways is a crucial glucose signal transduction pathway in plants [60].

For SnRK1/TOR signaling transduction pathway indirectly responds to glucose signaling by sensing energy consuming and environment stress in plant, which control the plant growth and development[61]. The Tre6P-aucrose transport between source organ such as leaf and sink organ such as seed, root. OsNAC02 as TF which involving in the DNA replication and cell cycle control, the variation of its expression could result into the SSC fluctuation. As these processing changed, the energy metabolism system changed either. The most obvious appearance is the floury endosperm formation at seed early development with 50% white-core, and the Ko-mutants are all premature as a result.

### Ko-mutants prematurity relative to chalky phenotype and quality change in the vegetative endosperm development

The seeds of all mutants mature about 15 days earlier than those of WT, and display the yellowish glumes in mutant and green caryopses in WT (95 DAG). Meanwhile, there is a great qualitative variation in the endosperm of mature and dry seeds in mutants vs WT. As the SSC is up-regulated significantly in the mutant seed (125DAG) (S7 Fig.), the reason of premature may be that the sugar is a key bio-signal for triggering the vegetative development, and a higher SSC in seed will enhance this signal including the sucrose, maltose, glucose, which regulated these processions on mRNA-level and post-mRNA-level by miR156 [62]. Integration of volcano plots and Mapman results with pro-pro String, the relative genes in the Starch and Sucrose metabolism pathway (Fig. 4D, Fig. 6B), such as *OsBAM2* (LOC_Os10g32810) and *OsSSIIIb* (LOC_Os04g53310) in the volcano-DEGs (S10 Fig.), the *RSUS1* (LOC_Os03g28330), *OsBE2b* (LOC_Os02g32660), *OsDPE2* (LOC_Os07g46790) in the Mapman KEGG, *GBSSII* (LOC_Os07g22930), *OsGLN2* (LOC_Os01g71670) in Venn analysis, which results primary changes in the quality of mutant seeds, the energy transformation from storage amyloplast to the chloroplast (**Fig. 7**, 8). SSC was upregulated and the DP ≤ 13 of amylopectin were obviously increased in the N2 mut, the chalkiness with white-core in mutant seeds induced through the amylopectin syntheses increasing, all of these phenotypes above have been reported previously[62–64]. These upregulated genes can induce chalky in endosperm and variations in physicochemical properties.

**Fig 8.**
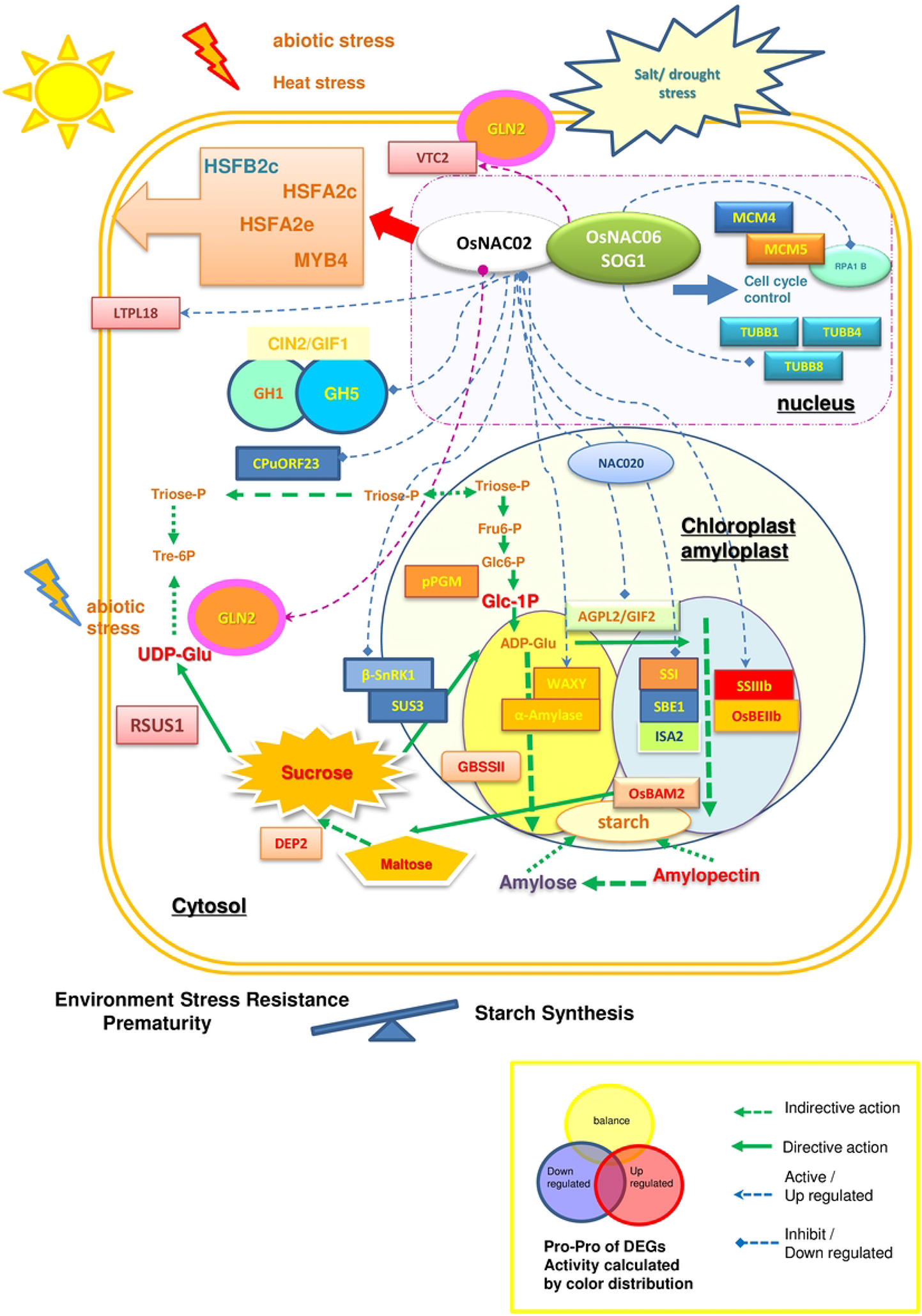
The antagonism between energy storage and development requirement under stress in Ko-OsNAC02 mutant. The pro-pro interactions in Ko-Osnac02 mutant are displayed by arrows with different shapes and different lines. The expressive activity of more than 30 proteins displayed by genes labeled with circles and rectangles in different colors. The blue is down-regulated while the red is up-regulated.

The *OsBE2b* (LOC_Os02g32660) overexpressed in N2 mutant which may have a slight increase in short amylopectin chains (DP=10-12), and this result is similar to that of an accumulation of excessive branched, water-soluble polysaccharides in OE-*BEIIb* mutant [65]. RSUS1 is presented in the aleurone of the developing seed and plays a role in sugar transport into endosperm cells [66, 67], but in the Ko-*Osnac02* mutant where the *RSUS3* was down-regulated in endosperm, which could increase the sucrose in seed. *OsBAM2* (LOC_Os10g32810) located in the chloroplast was up-regulated in N2 mutant, and it could increase the maltose content and inducing excessive amylopectin decreased into a balance[68]. Meanwhile, OsDPE2 (LOC_Os07g46790) responds to the maltose increasing stress and transport the maltose to other glycogen, in order to decrease the maltose ratio in seeds [69]. As the soluble sugar in endosperm increased, the juvenile to adult phase is shortened, and the resistance to abiotic stress is increased in mutant as well. *OsGLN* (LOC_Os01g71670) encode an enzyme endo-1,3-b-glucanase (1,3-b-GLU), which is expressed in the germinating seed [70]. It belonged to a huge family named glycosyl hydrolases family17 in plants, involving in the transportation of sugar onto a lipophilic acceptor which changed its chemical properties, and alters the bioactivity, enabling access to membrane transporter systems[71, 72]. The glycosylation of *OsGLN* is important for improving the resistance of abiotic stress such as salt stress in rice [73].

In the early stage of endosperm development, the amylose content in seeds (5 DAP) of N2 mutant was significantly lower compared to N3 mutant (S7F Fig.), which is contrary to the result that the amylose content in N2 mutant seeds (125 DAG) significantly decreased to a lower level at mature stage (Fig. 3B). This result may be induced by the increased expression of *OsSSIIIb* (LOC_Os04g53310) in the endosperm and *GBSSII* (LOC_Os07g22930) in the chloroplast, which means more energy was stored in leaf instead of being transported to the sucrose-sink in endosperm, and there were more amylopectin and less amylose in N2 seeds compared to WT and N3.

Combining the DEGs in pro-pro interaction and Mapman KEGG based on seeds transcriptome (Fig. 6B), all the key genes down-regulated are including *β-SnRK1* (LOC_Os05g41220), *SBEI* (LOC_Os06g51084), *OsSUS3*(LOC_Os07g42490). While *OsCIN1*(LOC_Os02g33110), *OsCIN2* (LOC_Os04g33740), *AGPL2* (LOC_Os01g44220), *ISA2* (LOC_Os05g32710) are expressed in no difference between N2/N3 and WT.

The *SBE I* (LOC_Os06g51084) are specially expressed in the filling grains, and mainly for synthesis of the medium and long amylopectin chains. The decreased expression of SBE I in Ko-*Osnac02* mutant may show a slight decrease in long amylopectin chains (DP > 37) and intermediate chains (DP=12-21), while a slight increase in short amylopectin chains (DP < 12) [74], but no significant change in the appearance and weight of the caryopsis are found as the results in Figure 3D [75, 76].

SnRK1 complex composed of α1, α2,β,γ subunits is the central metabolic regulator in the sucrose-dependent precession, and it is specific in the plant which is regulated by variations of sucrose content. *β-*SnRK1 (LOC_Os05g41220) together with γ-SnRK1 are two regulatory subunits of SnRK1 complex[77], this gene down-regulated in N2 *mut* can inhibit the sucrose transferring to ethanol and glycerol in cytoplast [78]. Meanwhile, the high content of sucrose stimulates T6P accumulation will inhibit the SnRK1 expression in the Suc-Tre6P nexus model [60, 79], this means that *SnRK1* down-regulated could promote the resources being used for reactivating growth and reproduction in order to balance high content of Tre6P (Fig. 6B). *OsSUS3* (LOC_Os07g42490) is named Rice Sucrose Synthase 3 for its expression in rice only, and predominantly in the seed endosperm where it played an important role in starch filling processes during the milky stage of rice seed ripening [80]. It has been selected as a key enzyme in the sucrose-to-starch conversion of rice spikelet during the grain filling under abiotic stress, and its overexpression enhances the thermo-tolerance in rice[51, 81]. As the first key enzyme in sucrose synthesis, the down-regulated expression of *OsSUS3* might be a feedback to the high sucrose content. Overall, all of these genes expressions in the starch synthesis in seeds (3 DAP) were downregulated during early endosperm development. This caused the smaller starch granules in the central of the endosperm, and the amylose and amylopectin contents of the mature seeds (125 DAG) were lower compared with WT.

### DEGs in Ko-mutants are relative to abiotic resistance in the vegetative endosperm

Integration of the volcano plot and Venn analysis in Figure 4 and the mapman KEGG in **Figure** 6B, and most of these genes are relative to enhancing abiotic stress resistance.

Such as endoglucanase 7 (LOC_Os02g50490), *OspPGM* (LOC_Os03g50480), *OsVTC2* (LOC_Os12g08810), *OsBAM2*(LOC_Os10g32810), *OsSSIIIb* (LOC_Os04g53310),*GBSSII* (LOC_Os07g22930), *LTPL18* (LOC_Os01g12020), *OsGLN2* (LOC_Os01g71670), *glutelin* (LOC_Os02g15070), *ycf68* domain (LOC_Os04g16722) are up-regulated in the DEGs. And most of these genes were relative to abiotic stress in N2 mutant.

Endoglucanase 7 (LOC_Os02g50490) was highly expressed in N2/N3 *mut*, and involved in cell wall synthesis and cell wall organization process under abiotic stress in roots[82, 83]. As these genes are highly up-regulated in N2 *mut*, which mainly promote the starch degradation and the soluble sucrose production, as well as enhance the cell resistance to abiotic stress. The *OspPGM* (LOC_Os03g50480) which significantly up-regulated in the N2 mainly related to the sterility of male pollination, and expressed highly at the starch synthesis stage [84]. This gene could respond to abiotic stress by sucrose-signal as well (Fig. 5B). *OsGLN2* which is primary expressed in the endosperms of germinating seeds where its expression is induced by GA and suppressed by ABA [70]. *OsVTC2* (LOC_Os12g08810) encodes the rate-limiting enzyme in the ascorbate synthesis pathway, and the plant with higher ascorbate content enhances the salt stress tolerance and Fe bioavailability [85]. These genes above are highly upregulated in N2 *mut*, and they mainly promote starch degradation and the soluble sucrose production, as well as enhance cell resistance to abiotic stress.

According to the down-regulated DEGs in the volcano (Fig. 4C and S12A Fig.), the most prominent genes are as follows: the *ONAC020* (LOC_Os01g01470), *GH1* (Os3BGlu6, LOC_Os03g11420), *GH5*(LOC_Os04g40510), *OsCPuORF23* (LOC_Os10g40550), *β-amylase* (LOC_Os03g22790), *OsTPP3* (LOC_Os07g43160). In this study, *GH1* (LOC_Os03g11420) and *GH5* (LOC_Os04g40510) as the *β-glucosidases* are expressed in the germinating seed or the leaves as cellulose tissues involving in carbohydrate metabolism in the vegetative development, which inhibit the seed filling rate. Their expression could be induced by the transient starch in KEGG pathways, and also be activated in the transient starch transportation [4, 86]. *CPuORF23* (LOC_Os10g40550) involved in trehalose synthesis, which is relative to drought resistance [42]. All these genes deletion can reduce the cell walls resistance to abiotic and biotic stress and delay the production of disaccharide as a substate transfer to cell walls synthesis.

### Formation of chalky seeds by the incomplete filling in the vegetative endosperm

According to the pro-pro co-network in seed (3DAP), three groups of co-networks genes were selected on the threshold (score ≥ 900), and seven genes of group I control the chalkiness formation in the early endosperm development, including: *Waxy* (LOC_Os06g04200), *OsAGPL2* (LOC_Os01g44220), *SSI* (LOC_Os06g06560), *SBE I* (LOC_Os06g51084), *ISA2* (LOC_Os05g32710) (Fig. 5B), besides with *ONAC020* (LOC_Os01g01470) (Fig. 4C).

In the whole starch metabolism pathway, *AGPL2* (LOC_Os01g44220), *SSI* (LOC_Os06g06560), *SBEI* (LOC_Os06g51084), *ISA2* (LOC_Os05g32710) are active in amylopectin synthesis, which are down-regulated in N2 seed (3DAP) mutant (Fig. 6C). Meanwhile, *Waxy* (LOC_Os06g04200) is active in amylose synthesis with a stable expression at normal-levels. In the previous study, knock-out or down-regulated these genes can induce chalkiness in rice endosperm.

As ADP-glucose pyrophosphorylase (AGP) large subunit that plays a crucial role in the regulation of starch biosynthesis and grain filling during rice seed development, the deletion of *OsAGPL2* and *OsAGPS2b* causes a shrunken endosperm due to a significant reduction in starch synthesis [87–90]. Through the yeast-two-hybrid assay, *OsAGPL2* interact with *OsAGPS1*, *OsAGPS2a* and *OsAGPS2b,* the expressions of granule-bound starch synthase, starch synthase, starch branching enzyme and starch debranching enzyme are dominantly decreased in *gif2* (OsAGPL2) mutant [91].

Waxy and dull rice grains have an opaque endosperm due to the pores between and within the starch granules[1]. *Waxy-I* bears a loss-of-function mutation that results in glutinous rice varieties with extremely low AC, *Waxy-II* shows a leaky phenotype with a moderate level of AC, and *Waxy III* functions as WT allele with high AC [92].

As homologous genes *ISA1* and *ISA2* have similar functions in the amylopectin precursor formation process in rice endosperm. FLO6 (CBM48) could interact with *ISA1* directly to participate in the relative starch protein folding process, and plays the same role as *SSG4* in controlling the size of starch granules, then the floury endosperm can be formed by controlling the size of the compound granules [93, 94].

In previous study of chalky rice, *OsNAC20* or *OsNAC26* caused starch and storage protein synthesis decreasing in their mutants. When the mutation of *OsNAC20* or *OsNAC26* occurs alone with on change in grains, but the *OsNAC20/26* double mutant could significantly decrease the starch and storage protein content. Because they could bind to the promoters of *AGPS2b*, *AGPL2*, and *SBEI*, and the expression of these three genes was severely decreased in the *OsNAC20/26* mutant [95]. As the *OsNAC020* (LOC_Os01g01470) is down-regulated in the early filling stage (Fig. 4C), then the storage of starch and protein decrease and the expression of *AGPL2*, *SBEI*, *Waxy* was consequently downregulated in the starch pathway (Fig. 6C).

Overview all the genes above in the starch synthesis in seeds (3DAP), the most gene expressions were down-regulated during the early development. This causes the starch granules smaller in the central of endosperm, and the content of amylose and amylopectin in mature seeds (125 DAG) was lower than that of WT. According to the pro-pro String data of interactive DEGs in RNA-seq based on the N2 *mut* seeds (3DAP) versus WT, the *OsSUS3* expression is down-regulated in the endosperm, however, the *RSUS1* in the elongated cell such as the aleurone is up-regulated. This may be the reason why both white-core and white-bell are formed in N2 seed, and this expression model may be an urgency strategy used for harsh environment resistance. Because the endosperm development was omitted while the aleurone, caryopses development of seeds in a normal rate, then the floury endosperm formed as a result, and this may be why the SSC content in N2 *mut* are much higher in mature seeds (125 DAG) compared with WT.

In conclusion, according to the qRT-PCR verification of Mapman KEGG in starch synthesis, 90% of genes in the ECs were down-regulated (Fig. 5B). The Starch is the main component of the rice embryo, which occupied 76.7 % - 78.4 % of fresh seed and 95 % weight in dry seed, while the storage proteins occupy only 8 % of the dried rice embryo. The different ratios of amylose and amylopectin combinations contribute the different quantities of rice embryos. In the early endosperm development, due to insufficient starch synthesis, the soluble substrates content such as amylose and soluble sugar was higher for the necessary energy up-conversion, followed by a chalky phenotype with a white-core and white-belly in the endosperm.

### Premature caryopses in N2 *mut* with cell cycle shorting in dividing cells

Cell cycle controlling genes such as the MCM family, Tublin proteins, NAC family are mainly expressed in dividing tissues, and their subcellular locations are mostly in the nucleus. Their expressions change with the DNA replication, and they are conserved in the domains of the protein structure in different plants.

The MCM contains six subunits and its substrate is poly AAA+ ATPase (Adenylate phosphorylase). After that, it can be anchored to a binding site on the DNA. MCM can interact with DNA to depolymerize, which controls one mitosis, and the stain volume of the DNA double-strand can be completed per cell cycle [96].

The MCM family of proteins play an important role in the chromosome replication process in eukaryotes [52], and MCM2-MCM7 are highly conserved in organisms. In the process of chromosome replication, the MCM DNA protein plays a role in stabilizing chromosomes during the cell cycle G0-G1, which is responsible for the precise replication of the chromosomal DNA. At the same time, the MCM is involved in the formation and stabilization of centrioles during mitosis. It was demonstrated that when the MCM expression decreased in the G0-G1 phase, it would lead to DNA damage and chromosomal abnormalities, activate the expression of ATR/ATM gene and induce expression of CHK1/CHK2 [54].

MCM4 and MCM5 play an important role in regulation of cell cycle from G0 to G1-phase. MCM7 is induced by E2F-1 and expressed in the late G1-phase, and it hardly expressed in the G0 phase, but its expression gradually increases during the transition from the G0 to the G1-phase. The loss of MCM7 in the Metaphase (G1,16 h) increases the probability of chromosome mismatch and abnormality, which will shorten the cell division cycle and premature the formation of abnormal chromosome. Meanwhile, defective or abnormal expression of MCM7 in the S-phase of mitosis will double the synaptic distance between the centriole and sister chromatids, leading the abnormal growth and development and induce cancer [54].

In animals, the absence of MCM7 induces DNA damage and chromosomal abnormalities, and any reduction in the expression of MCM4 also lead to DNA abnormalities. In this research, we found that *MCM4* (LOC_Os01g36390) showed a low-expression in N2 *mut*, while the *MCM5* (LOC_Os02g55410) and *MCM7* (LOC_Os12g37400) expressed at normal levels. The proposed conclusions on the shortage of grain filling period: First, the *MCM4* is directly regulated by *OsNAC02* for low expression in the Ko-*Osnac02* mutant. *MCM4* as the core subunit of the MCM2-MCM7 complex in animals, and the deletion of *MCM4* may form a special minichromosome-loss phenotype which can induced apoptosis and death with homozygosity by hromosomal abnormalities [97]. This function is the same in plants. Second, MCM plays an important role in stabilizing chromosomes during chromosome replication and division. Deletion induces a DNA damage stress response that was previously due to cellular mechanical damage. It was report that the endogenous MCM5 and MCM7 proteins are equivalently localized in the nucleus during the G1, S, and G2 phases of the cell cycle and are released into the cytoplasmic compartment during mitosis[53]. As the *MCM5* (LOC_Os02g55410) and *MCM7* (LOC_Os12g37400) expression were up-regulated in N2 mutant. Although previous reported that MCM overexpression induces cancer in animals, it has not been proven that *MCM2* is overexpressed in plant culture tissues with normal DNA replication, which means that a moderate over-expression of MCM shortens the cell cycle, thus inducing earlier seed maturation.

## Conclusion

In the OsNAC02 co-network based on DEGs of N2 versus WT (**Fig. 8**), the effective genes regulated by OsNAC02 are mainly TFs, and they respond to both heat stress and other abiotic stress for the multiple functional motifs in their promoter regions. OsNAC06 is mainly involved in regulating cell cycle changes and speeding up endosperm filling rates under the environmental stress; all of these processes are controlled by a series of genes expressed in the nucleus, which determine chromosome replication. The down-regulated expression of these genes, such as MCMs and TUBBs, accelerates the replication cycle and rate, which also affects energy storage and speed consumption.

From the DEGs in the volcano plot (Fig. 4D), the Mapman overview on Sucrose and Starch metabolic pathway (Fig. 6), the pro-pro co-network (**Fig. 7**), 50% of the crucial genes in ECs are collected by statistical analysis (log2 FC, p < 0.05). The important enzymes in the pathway are including: BAM2, SSIIIb, GBSSII, RSUS1, pPGM were upregulated, as well as CIN2(GIF1), AGPL2(GIF2), WAXY, ISA2 are expressed with no significance, and SBEI was downregulated in N2 mut. The regulation of these genes promoted micromolecular production such as sucrose, whereas the macromolecular storage of substrates decreased during the early seed-filling stage. All of the results above could cause floury endosperm in mutants. It is confirmed that the decreased AC in the N2 mut endosperm, as well as the SSC increasing significantly in N2 mut seeds (125 DAG) are relative to the upregulated expression of *RSUS1*(LOC_Os03g28330), *OspPGM* (LOC_Os03g50480) (Fig. 6B) and *OsBAM2* (LOC_Os10g32810) in the DEGs of N2/N3 mutant versus WT (Fig. 4C).

In the *Ko-Osnac02* mutant, a decreased energy storage substrate induced the soluble sucrose content increasing, which promotes faster development and enhances the resistance to the biotic and abiotic stresses in the whole plant.

## Data and protocols availability

The raw data of rice genome as the reference or control in RNA-Seq have been deposited in the Genome Sequence Archive (Genomics, Proteomics & Bioinformatics, 2021). The raw data of RNA-Seq in this manuscript have been deposited in the National Genomics Data Center (https://ngdc.cncb.ac.cn/gsa/, Nucleic Acids Res 2022), the China National Center for Bioinformation Beijing Institute of Genomics, Chinese Academy of Sciences (GSA: CRA008460), which are publicly accessible online (https://ngdc.cncb.ac.cn/gsa/browse/CRA008460). The detections of physicochemical properties are according to the procedures in our lab, and you could contact Dr. Wei for the details of the detection methods.

## Accession numbers

Gene Bank: OsNAC02, LOC4329852, OSNPB_020594800, LOC_Os02g38130;

OsNAC06, LOC4340715, OSNPB_060267500.

RAP_Locus : OsNAC06(Os02g020594800) OsNAC06(Os06g0267500);

Sequence data from this article can be found in the GenBank/EMBL, Processed RNA-seq data can be accessed and visualized at NCBI, and the platform website: http://www.lc-bio.com/

The gene ID in RNA-seq is accordance with MSU_loc (Rice Genome Annotation Project (uga.edu)) OsNAC02(LOC_Os02g38130), OsMYB04(LOC_Os04g43680), OsHSFA2E (LOC_Os03g58160), OsHSFA2C(LOC_Os10g28340), OsHSFB2c(LOC_Os09g35790);

Glutelin (LOC_Os02g15070), α-amylase (LOC_Os06g49970), OsBAM2(LOC_Os10g32810); SSI(LOC_Os06g06560), OsVTC2 (LOC_Os12g08810), endoglucanase7 (LOC_Os02g50490), RSUS1 (LOC_Os03g28330), OspPGM (LOC_Os03g50480), OsBE2b (LOC_Os02g32660), OsDPE2 (LOC_Os07g46790), OsGLN2 (LOC_Os01g71670), glutelin (LOC_Os02g15070), ycf68 domain (LOC_Os04g16722), OsSSIIIb (LOC_Os04g53310), GBSSII (LOC_Os07g22930), LTPL18 (LOC_Os01g12020);

MCM4 (LOC_Os01g36390), MCM5(LOC_Os02g55410), MCM7(LOC_Os12g37400), OsTUB1 (LOC_Os01g18050), OsTUB4 (LOC_Os01g59150), OsTUB8(LOC_Os03g45920), RPA1B(LOC_Os03g11540);

β-SnRK1 (LOC_Os05g41220), SBE1 (LOC_Os06g51084),CIN2(LOC_Os04g33740), AGPL2 (LOC_Os01g44220), ISA2(LOC_Os05g32710), OsSUS3 (LOC_Os07g42490);

OsNAC20 (LOC_Os01g01470), OsPUP2 (LOC_Os09g29210),OsTPP3 (LOC_Os07g43160), BBTI7 (LOC_Os01g03390), Aspartokinase (LOC_Os03g63330), Hsp20(LOC_Os04g36750), CPuORF23 (LOC_Os10g40550);

Os3BGlu6 (LOC_Os03g11420), GH5 (LOC_Os04g40510), ZmEBE-1 (LOC_Os01g26320),

GDSL like synthase (LOC_Os01g46220), CSLA9 (LOC_Os06g42020).

The qRT-PCR verification of 39 genes in 29 ECs in Starch and Sucrose metabolic pathways are selected from DEGs of mapman analysis in RNA-Seq, and the Accession Numbers in details are listed in Table S12.

## Supporting Information

S1 Fig. The mutant detection by PCR sequencing

S2 Fig. The yield characters of T2 N2 /N3 mut (95DAG) in field (95DAG in 2021, 125DAG in 2020)

S3 Fig. The homologous clusters of OsNAC family genes by phylogenetic analysis

S4 Fig. The homologous analysis of NAC in plant

The Gene ID are according to the database RAP (rapdb.dna.affrc.go.jp), and (OsNAC02, Os02g0594800), (OsNAC06, Os06g0267500) in NCBI of homologous analysis.

S5 Fig. The ABRE/ G-box motif locations on the promoter regions of OsNAC02 regulated genes

S6 Fig. The chalky phenotype of N2/N3 *mut* seeds (T2) in 2021

S7 Fig. The histological analysis during endosperm development

S8 Fig. The PCA clusters based on RNA-seq analysis

S9 Fig. The GO enrichment of DEGs in N2/N3 mutant vs WT

S10 Fig. The DEGs clusters in the Volcano plots of RNA-seq in seeds (3DAP).

S11 Fig. The genes of 39 ECs in Sucrose and Starch Metabolism pathway in Mapman of N2 vs WT seeds (3DAP) are verified by qRT-PCR.

S12 Fig. The model of starch synthesis in amyloplast /chloroplast of N2 mut

Table S1 The H-1-Y vectors construction of genes in OsNAC02 co-network.xlsx

Table S2 The data of WTVSN2_YVSN2_Z_Gene_differential_expression notation.xlsx

Table S3 The data of WTVSN3_1VSN3_3_Gene_differential_expression notation.xlsx

Table S4 The GO enrichment analysis of N2 vs WT and N3 vs WT respectively.xlsx

Table S5 The genes co-expressed in the down-regulated Venn diagram

Table S6 The up- and down-regulated genes notation of Venn diagram in seed (3DAP).xlsx

Table S7 The data of the WTVSN2_XVSN3_Y DEGs in Volcano diagram.xlsx

Table S8 The volcano diagram of genes labeled in DEGs of the seeds

Table S9 The rawdata DEGs in volcano diagram of Starch and Sucrose pathway.xlsx

Table S10 The volcano plot of genes labeded in Starch and Sucrose pathway of the seeds

Table S11 The two pro-pro interaction notations of WTVSN2_xVSN2_y _DEGs in heatmaps.xlsx

Table S12 The qRT-PCR primers of DEGs in starch pathway ECs for Mapman verification.xlsx

Table S13 The qRT-PCR primers of DEGs in co-network of OsNAC02.xlsx

## ACKNOLEDGEMENTS

The author would thank Dr Jiao providing the technology service for the physicochemical detection in milled rice and the Thank Dr. Chen give valuable suggestion for the standards of data analysis, Dr. Shao provide support to make sure all the experiment completed in available way. Thanks to the Pro. Hu and Pro.Tang, and the young managers of this key lab platform including X. J.,Wei, S.K.,Hu, Z.H.,Sheng, who provide the great environment for the experiment processing in CNRRI.

---

**S1 Fig.**
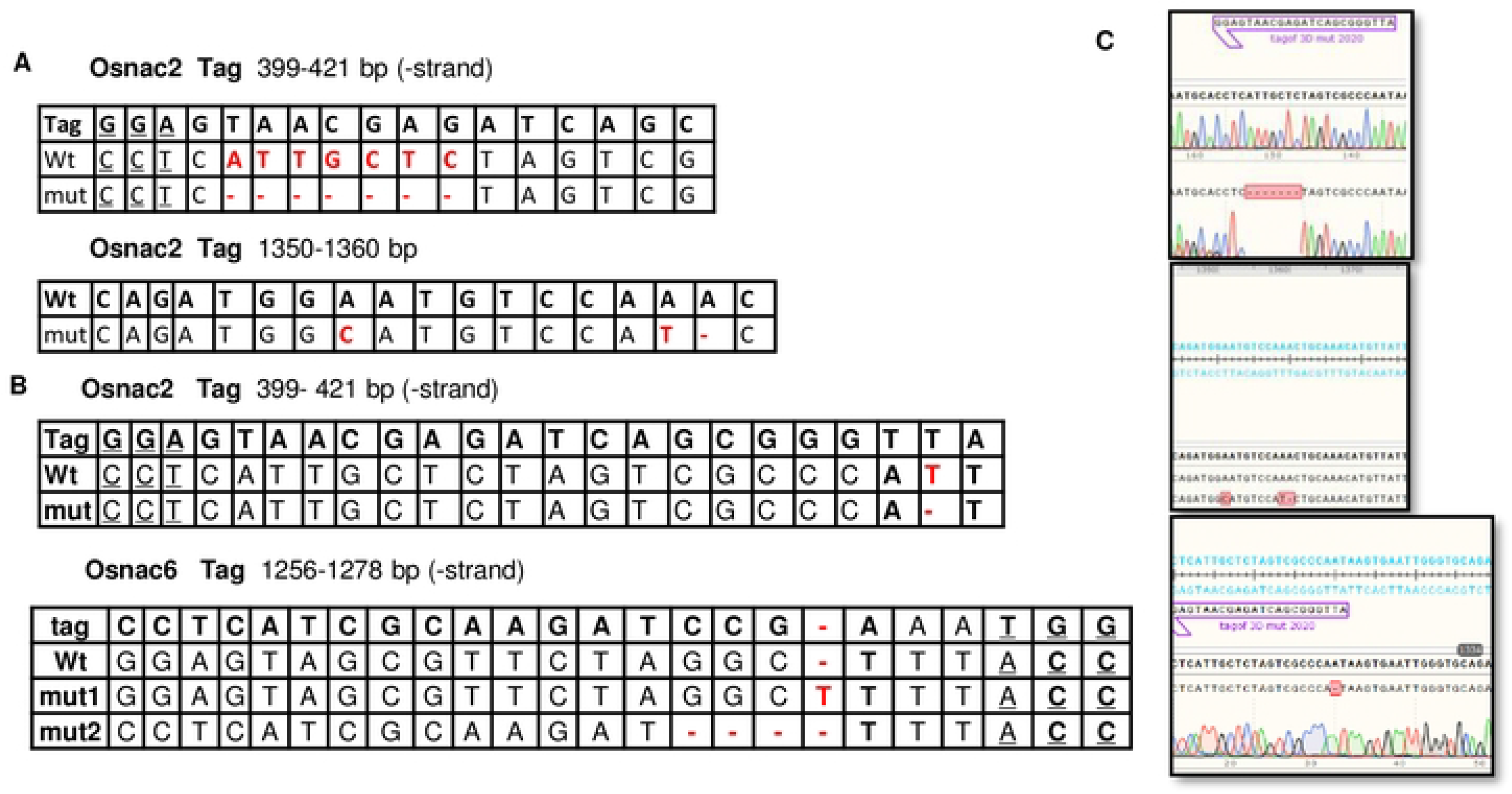
The mutants detection by PCR sequencing. **(A)** Ko-Osnac02 mut construction with two mutant sites in OsNAC02 CDS. The genetic type of mutants are labeled in red and the PAM “GGA” is highlighted in bold and underline. The mutant with “ATTGCTC” 7 bp-deletion in N2 mut located at 399-421 bp of OsNAC02 sequence, and the mutant with “A-C” at 1350 bp, A-T “ I” A-x “transversion at 1360 bp of OsNAC06 sequence. **(B)** N3 mut constructed the mutant sites in Osnac02 / Osnac06 CDS with PAM “GGA”/ “TGG” respectively. The Crispr-Cas9 induced “T” deletion in N2 and “T” insertion and 3 bp-“GGC” deletion in N3 at 1256-1278bp region respectively. **(C)** As these figures shows, the mutant sites of N2/N3 mut in OsNAC02 CDS detected by the alignment with DNA sequence of WT (NIP).

**S2 Fig.**
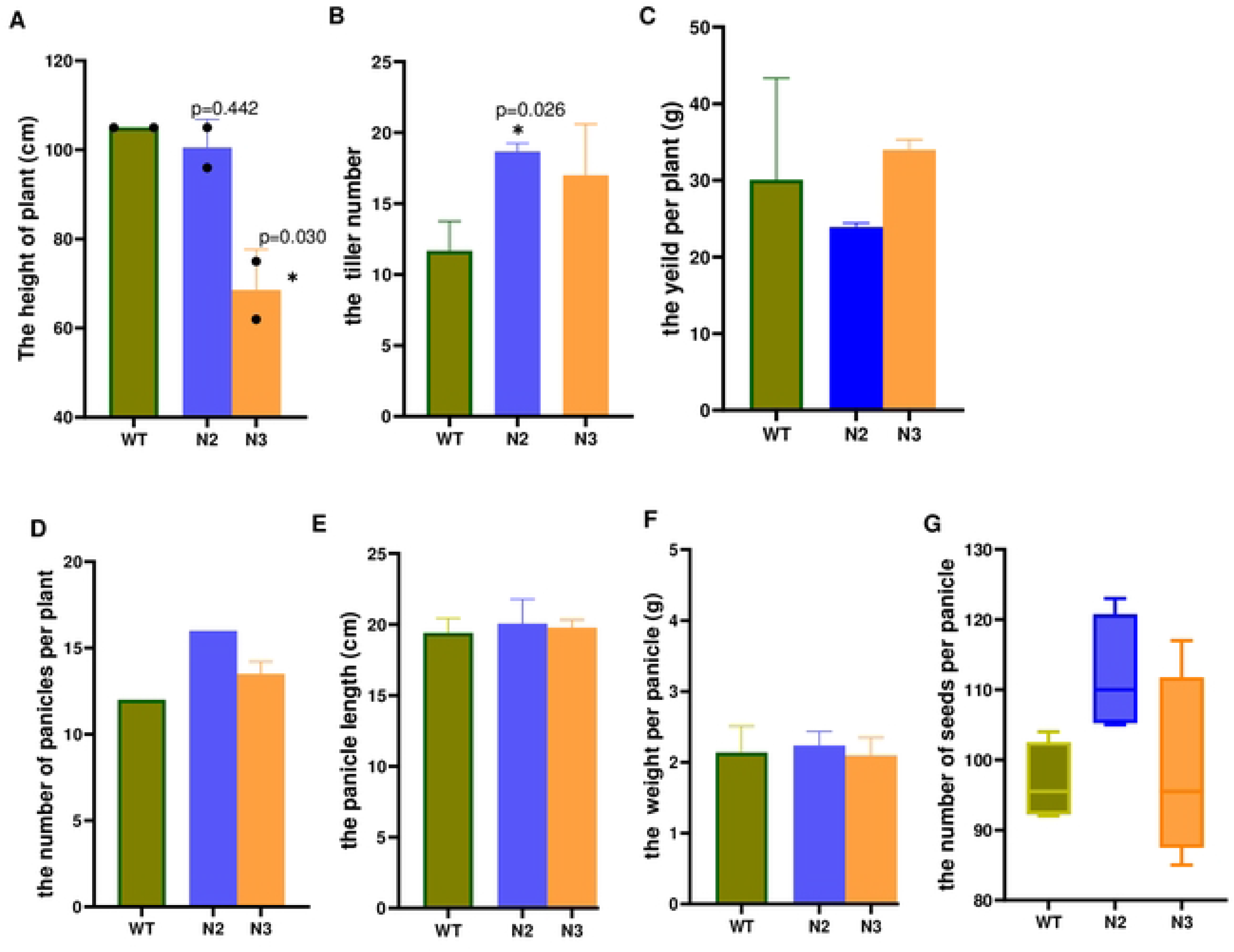
The yield characters of N2 / N3 mut in field (95DAG in 2021, 125DAG in 2020) **(A)** The plant height of N2/N3 mutant (T1) shows obvious difference in 2021, and both of them were dwarf compared with the WT. The height of N3 is higher than N2 mutant. **(B)** The number of tillers in N2 and N3 mutant are more compared with WT in 2021. **(C)** The N2/N3 mutant (TO) harvested in Hainan are detected for the yield per plant. All the seeds of T1 generation (125 DAG) are pre-treated with drying in the green house for two months. The yield per plant in N3 mut was heaviest and N2 mut was the lightest as a result. **(D)** The number of panicles per plant (950AG) in N2/N3 mutant, and the plant height, the tiller number per plant are detected simultaneously in 2020. **(E)** The panicle length per plant (95DAG) in N2/N3 mutant was detected in 2021. **(F)** The weight per panicles (95DAG) of N2/N3 mutant are detected, and there is no significant difference between mutants and WT. Five panicles from each plant are selected randomly for detection respectively. **(G)** The number of seeds per panicle in T1 generation (95DAG) was detected, and N2 mut has most seeds per plant. This phenotype together was detected with the number of panicles together in the same materials. All the mutants planted in field under the normal conditions (Fuyang, Hangzhou in 2021),the statistic analysis are running by the **student** t-test,*, p < 0.05 and **, p < 0.01. The T0 generation of transplants was planted in Lingshui (Hainan Province Island, 18.5°11O’E) in the winter of 2020, and we obtained the T1 generation. Following the T1 generation was planted in Fuyang (Hangzhou in Zhejiang Province, 30°120’E) during the summer in 2021, and T2 generation were harvested in the autumn of 2022. As the yield characters shown in **Figure 1**, the shape phenotype in seeds (125 DAG) of N2/N3 mutant are no difference compared with the control (nip).

**S3 Fig.**
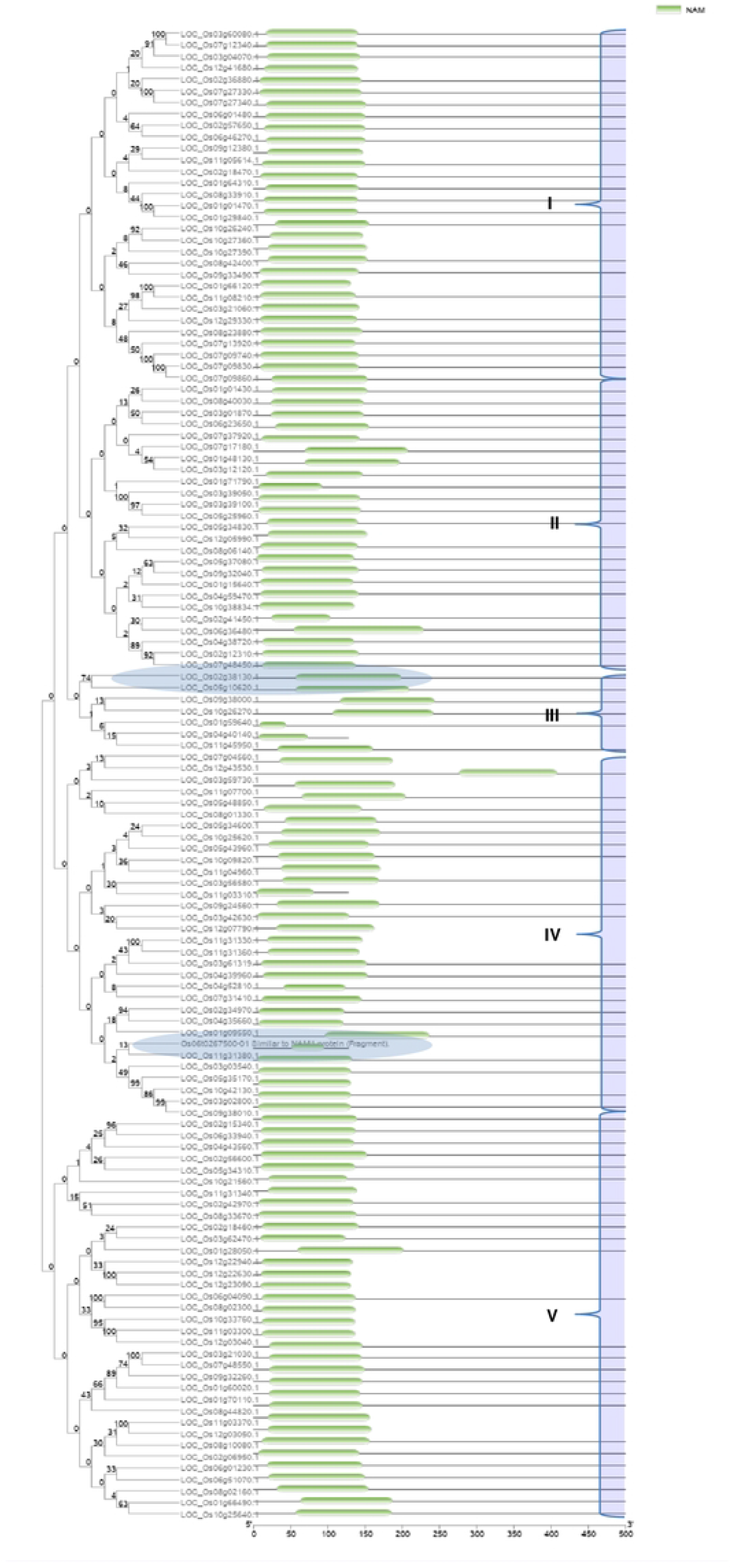
The homologous clusters of OsNAC family genes by phylogenetic analysis. The OsNAC02 (Os02g0594800, LOC_Os02g38130) and OsNAC06 (Os06g0267500)are labeled by bule circle. The two genes are belonged to subgroup Ill and IV respectively. The phylogenetic tree of OsNAC family was constructed using the MEGA 7.0. and the NAM regions in all genes are highlighted by TBtools11.2. The green boxes in the phylogenetic tree are NAC domains of genes.

**S4 Fig.**
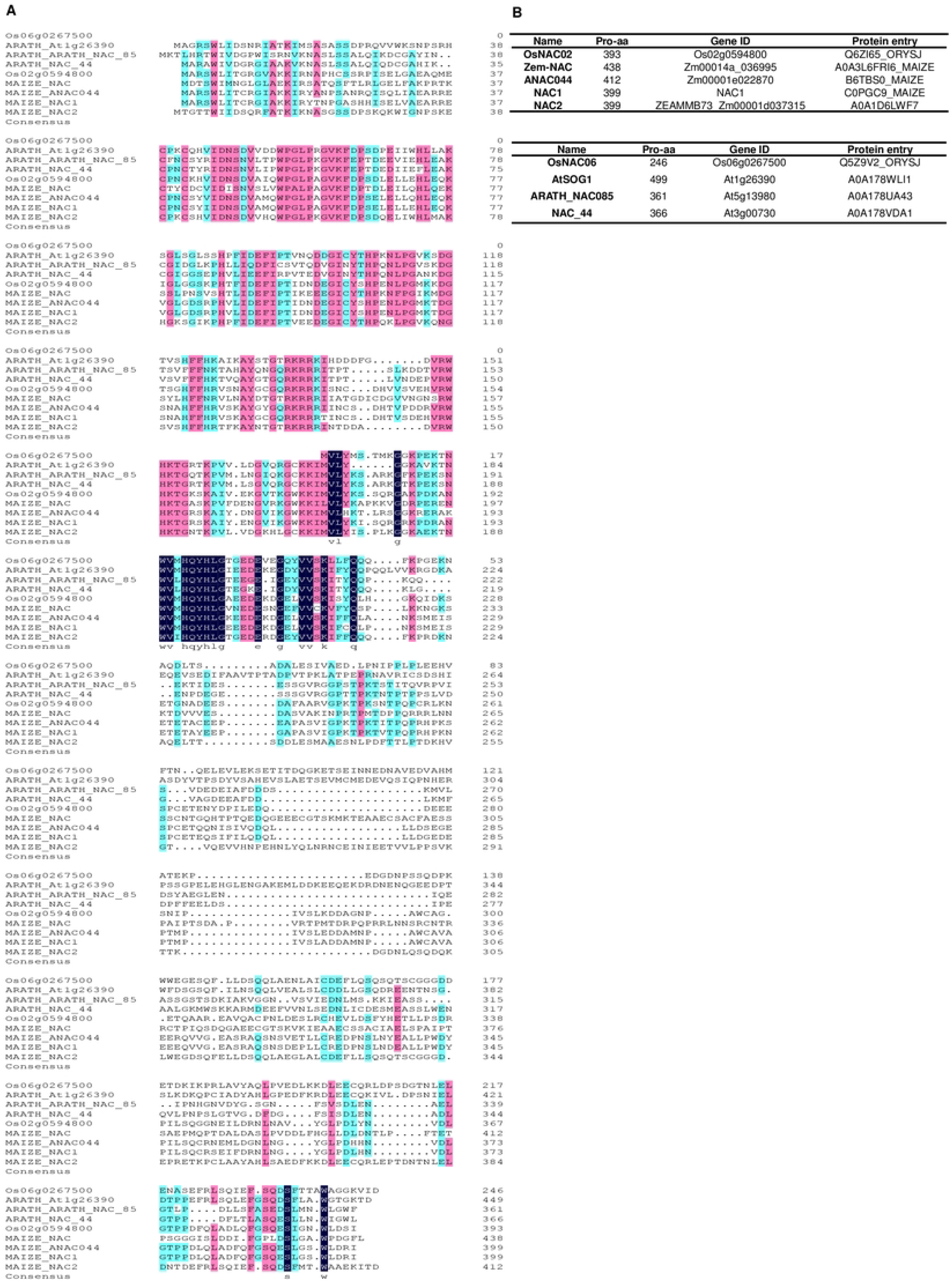
The homologous analysis of NAC in plant. **(A)** The homologous analysis of NAC proteins with amino acid sequence alignment in the rice, maize and Arabidopsis by using DNAMAN 7.0. **(B)** The Uniprot accession of 9 genes in homologous analysis. (https://www.uniprot.org/). OsNAC06 (Os06g0267500) has simple protein constructure and contains only 246 aa, which was highly homologous to AtSOG1. Its N­ terminal region with O - 45aa was a strictly conserved region of NAM domain.

**S5 Fig.**
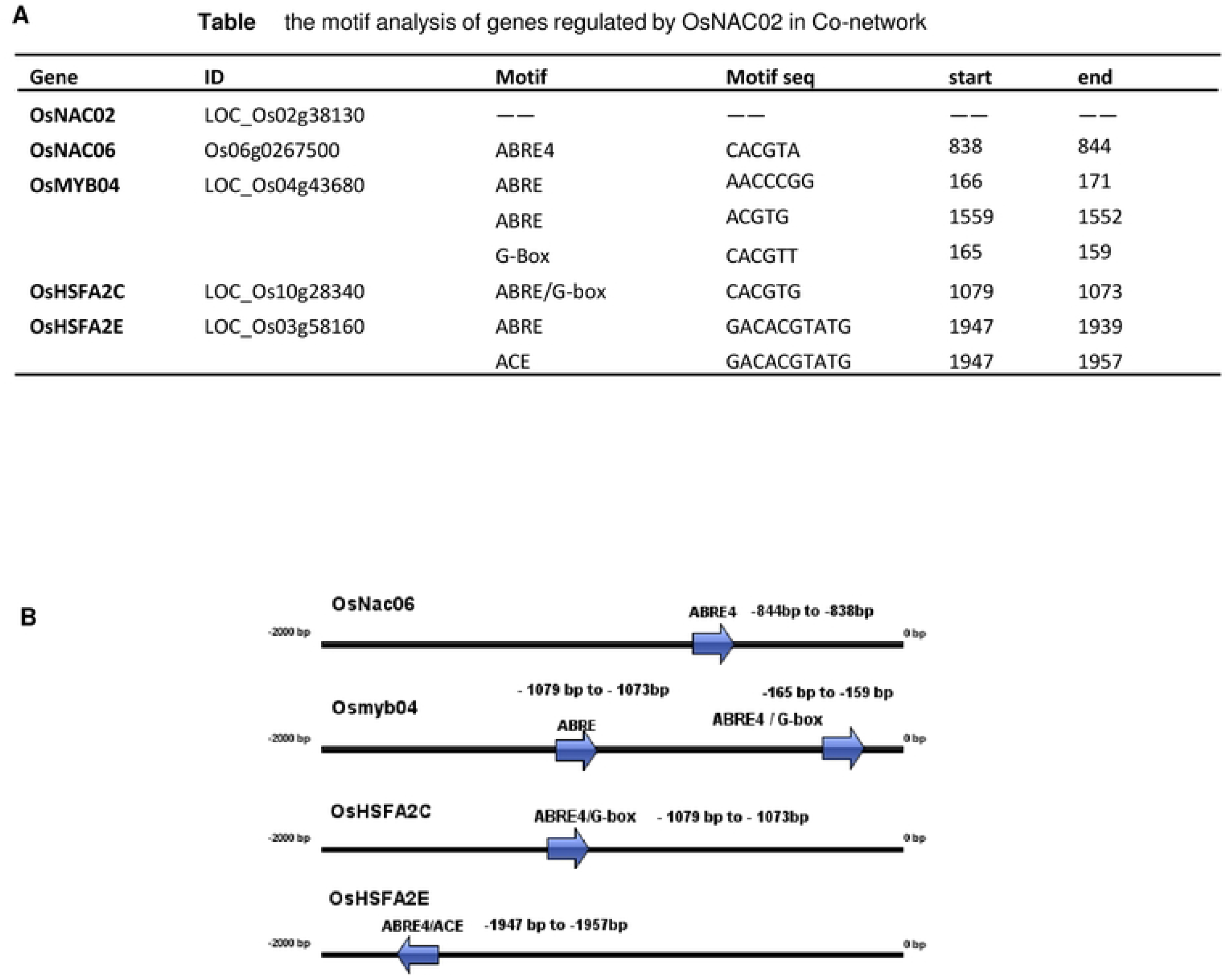
The ABRE/ G-box motif locations on the promoter regions of OsNAC02 regulated genes. **(A)** The motif analysis of genes regulated by OsNAC02 in co-network, all the location are detected by the PlantCARE. (http://bioinformatics.psb.ugent.be/webtools/plantcare). **(B)** The model of the motif analysis in the promoter region (-2000bp•’ATG’) of the OsNAC02 effective genes, as the arrow shows the direction of motifs with(+/-) on the promoter and the fig construction was completed by the software 18S1.0.

**S6 Fig.**
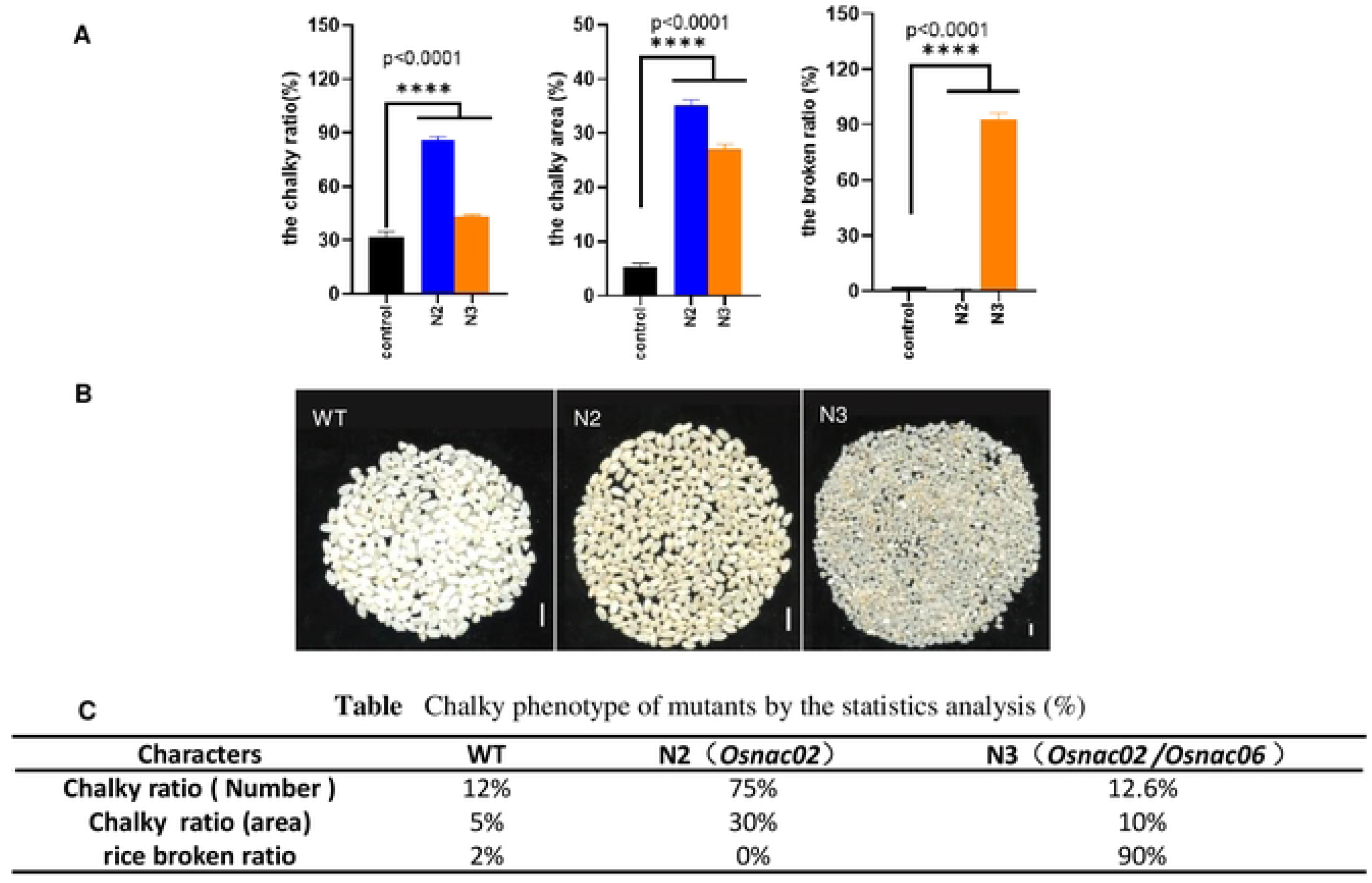
The chalky phenotype of N2/N3 mut seeds (T2) in 2021. **(A)** The chalky phenotypes of the milled rice are calculated by chalky ratio and the chalky area. The transparent are low in both mutant and WT, but the chalky ratio in mutants are obvious higher than that in WT. The rice broken ratio of N3 mut is highest of all. All the statistic analysis were performed by running a two-way ANOVA on the mixed model, *, p < 0.05, **, p < 0.01 and ***, p<0.0001. **(B)** The sample names are WT, N2 and N3 in the figure respectively. As the chalky phenotypes of milled rice and broken ratio of N2 and N3 mut per plant shown in this figure, both N2 and N3 are core-white in milled rice, but only N2 has core and belly white in its floury endosperm. As these figures show the yield per plant simultaneously, only N3 has highest yield and broken ratio in the milled rice per plant. n=10 mm. **(C)** The three phenotypes of transparent in N2 and N3 mutant by statistics analysis.

**S7 Fig.**
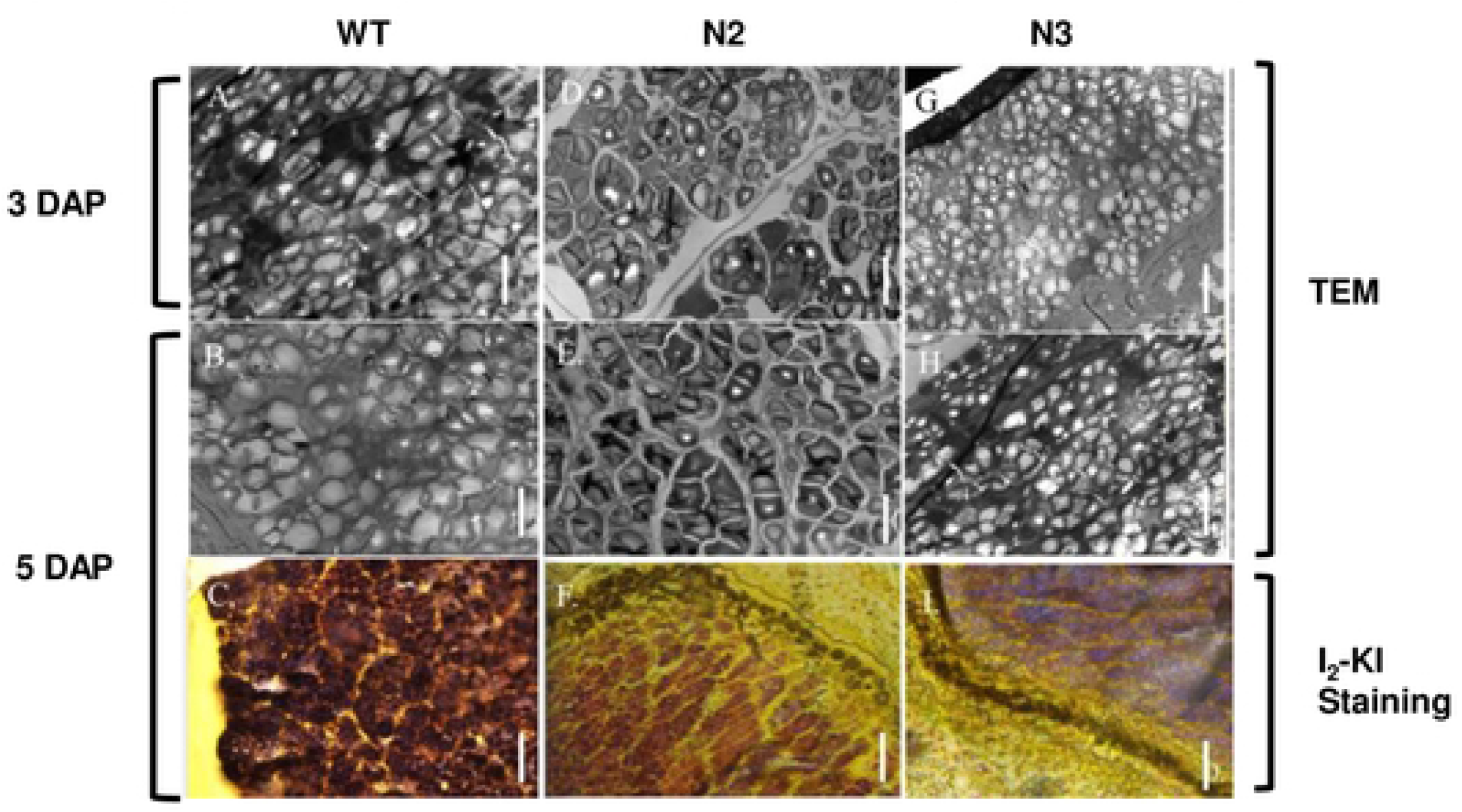
The histological analysis during endosperm development. Histological analysis of endosperm for maturation detection. As above shows the semi-thin sections of the opaque endosperm scanning in TEM and Kl-12 stains. n=100 um. The processes in endosperm development of WT/N2/N3 are shown in this figure respectively. As the **fig C**, **fig F**, **fig L** show the semi-thin sections stained of opaque endosperms at 5 DAP by the I_2_-KI.The iodine-staining(**l_2_-KI**) semi-sections implied the distinct starch content between N3 mut and N2 mut. For a darker blue in N3 mut (5 DAP) implies there are higher Amylase content in N3 mut (**fig L**) than that in N2 mut (**fig F**). **Abrreviation:** TEM, transmission electron microscope; OAP, days after Pollination.

**S8 Fig.**
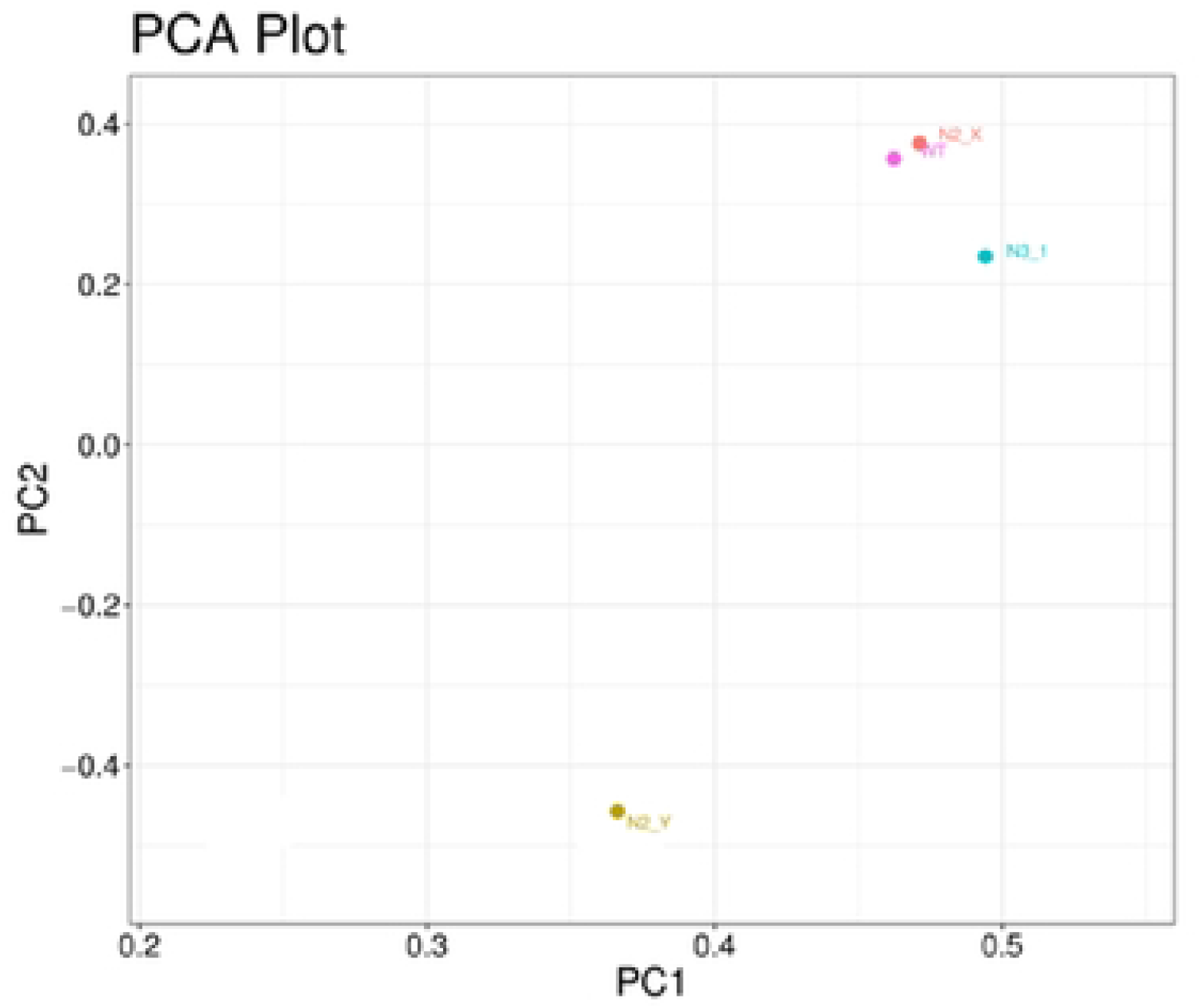
PCA clusters based on RNA-seq analysis. The trend of PCA analysis are based on the RNA-seq analysis of seed (3 DAP),and four samples are clustered into 3 PCA groups: The red spot was PCA1 cluster with WT and N2-1 (N2-x); The yellow spot was PCA2 cluster with N2-2 (N2-y), which used as duplicate sample of N2-x for the KO-OsNAC02 mut; The blue spot was PCA3 cluster with N3_1 only.

**S9 Fig.**
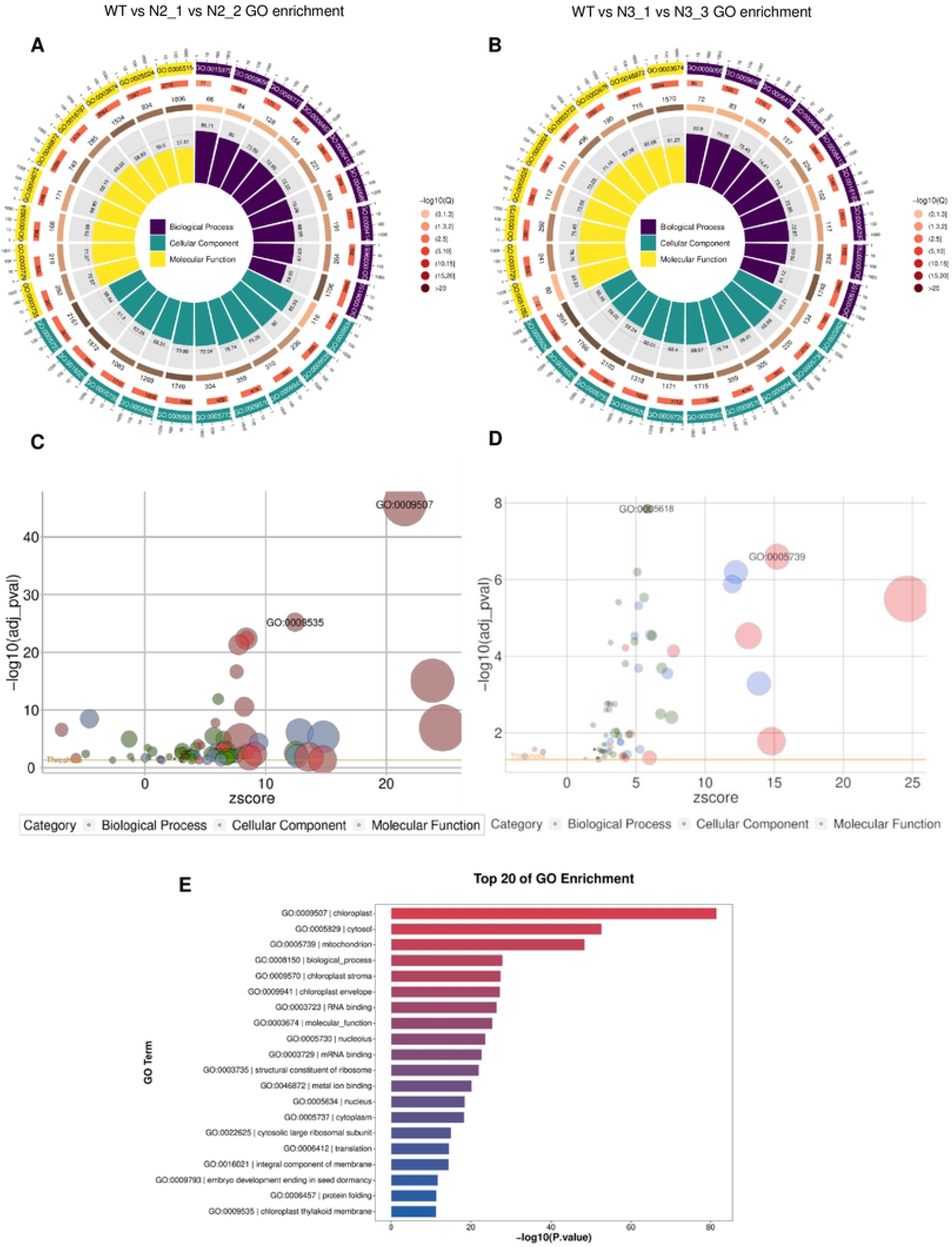
The GO enrichment of DEGs in N2/ N3 mutant vs WT. The gene enrichments of the GO analysis including 3 indexes: the top 9 GO pathways of the biological process (purple), the cellular component (green), the molecular function (yellow). **(A)** The top 9 pathways of GO enrichment in **N2 vs WT** **(B)** The top 9 pathways of GO enrichment in **N3 vs WT**. **(C)** The GO bubble Statistics includes biological process (blue),cellular component (red), molecular function (green), which were in accordance with results in **fig A** and **fig E**. In thecellular component (red), the two GO pathways :GO :0009507-thechloroplast pathway and G0:0009535-the chloroplast thylakoid membrane pathway were significantly enriched in **N2 vs WT**. **(D)** The bubble analysis of Go pathways in the **N3 vs WT**. In molecular function (green), the G0:0005618-cell wall pathway is the most significantly enriched pathway. While less genes are enriched in the other cellular component (red) G0:0005739-mitochondrion pathway. **(E)** The results represents the top 20 GO pathways of seed transcriptome including the GO significantly enriched pathways as appearing in **fig C** and **fig D**.

**Figure S10.**
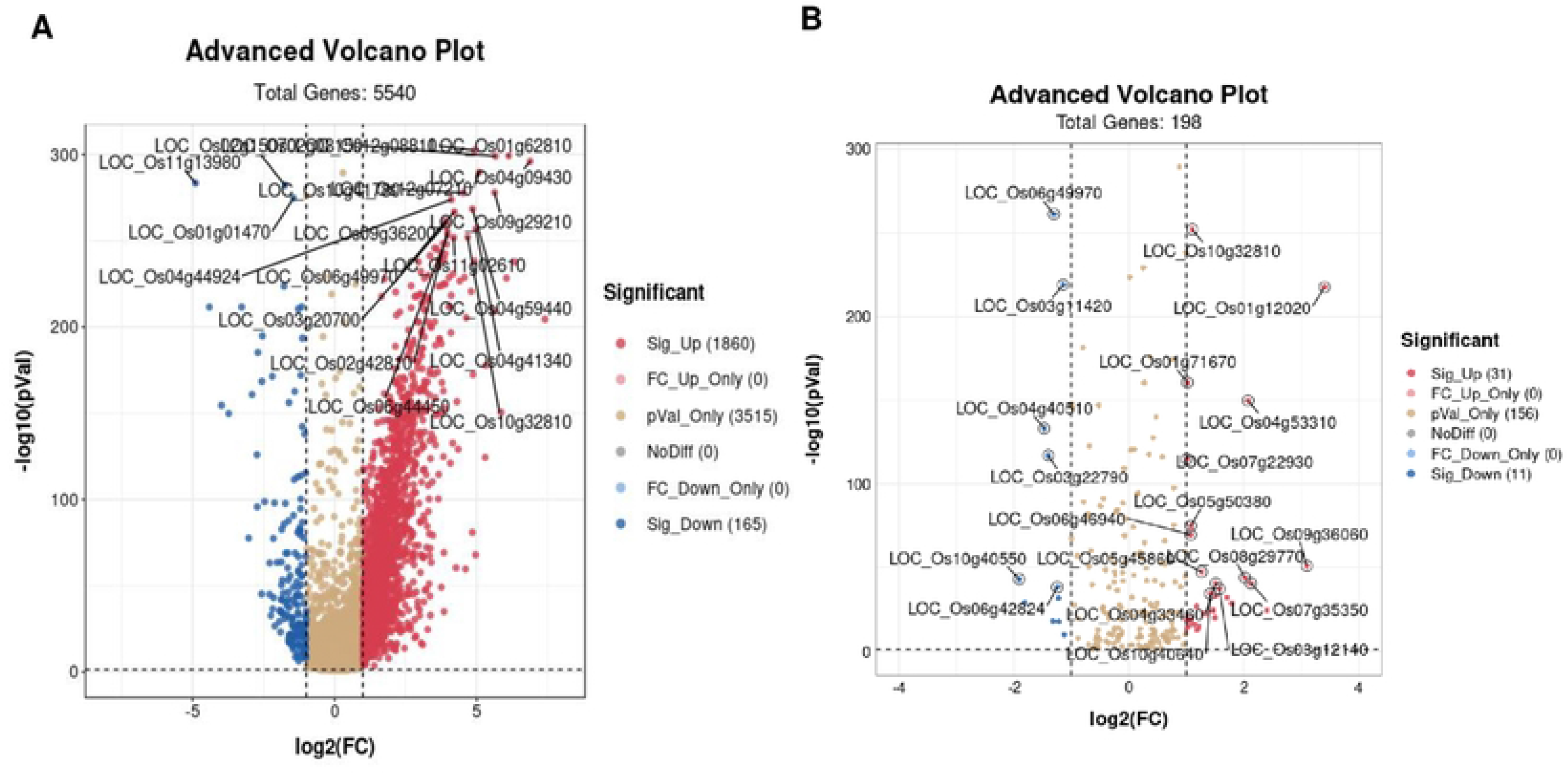
The DEGs clusters in the Volcano plots of RNA-seq in seeds (3DAP) The top 20 genes in DEGs are labeled with loc_lDs in the Volcano plots. **(A)** Volcano analysis showed the distinctive expressed genes (DEGs) in seeds of mutants vs WT, and the most genes of DEGs are up-regulated. **(B)** Volcano analysis showed the DEGs of all the relevant genes in the Starch and Source metabolism pathway in the seeds of mutants vs WT.

**S11 Fig.**
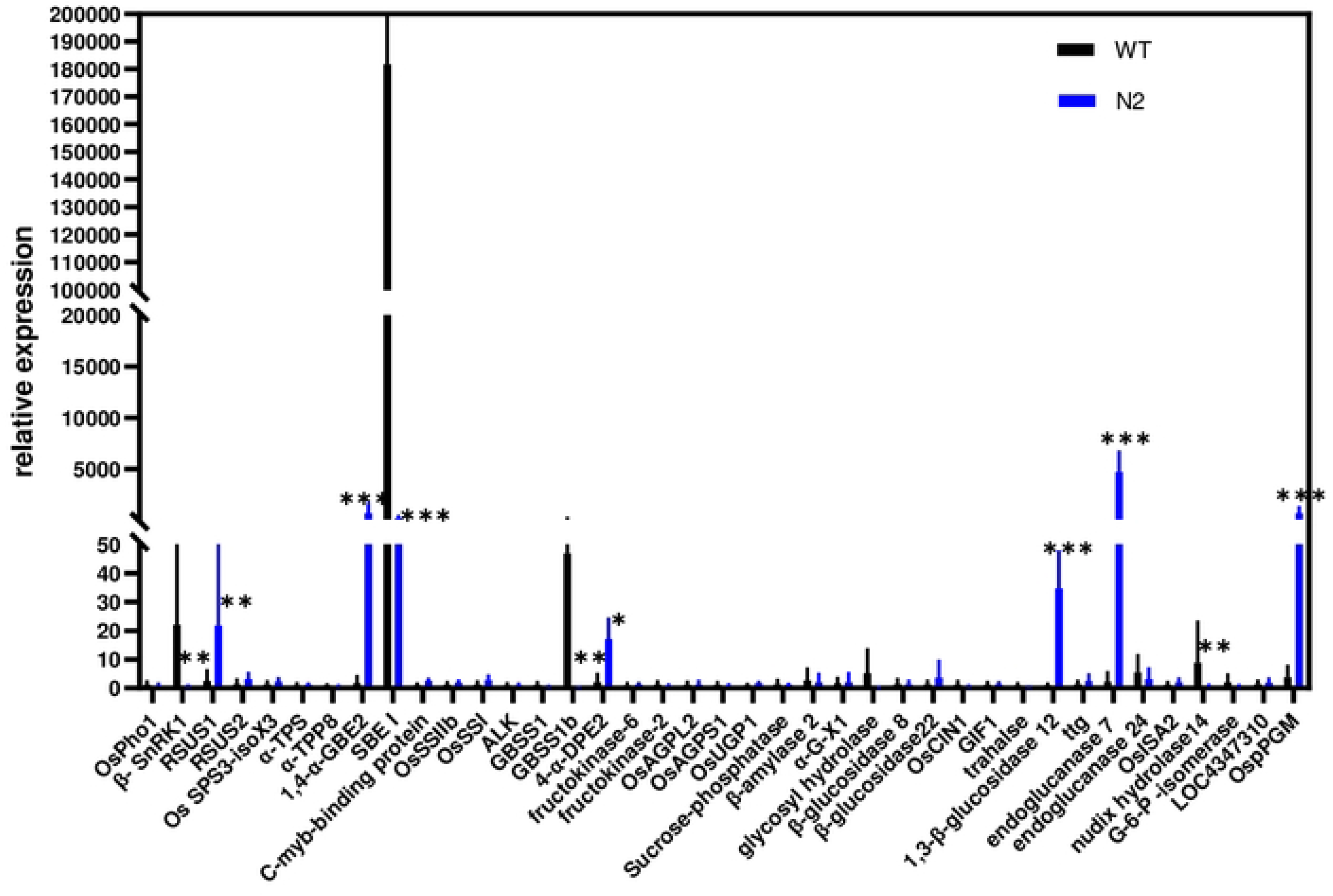
The genes of 39 ECs in Sucrose and Starch Metabolism pathway in Mapman of N2 vs WT seeds (3DAP) are verified by qRT-PCR. The qRT-PCR verification in seeds (3DAP) displayed the expressions of OsNAC02 relative genes in its regulatory co-network. All the statistical analyses are performed by student I-test, *, p < 0.05, **, p < 0.01 and ***, p < 0.001.

**Figure S12.**
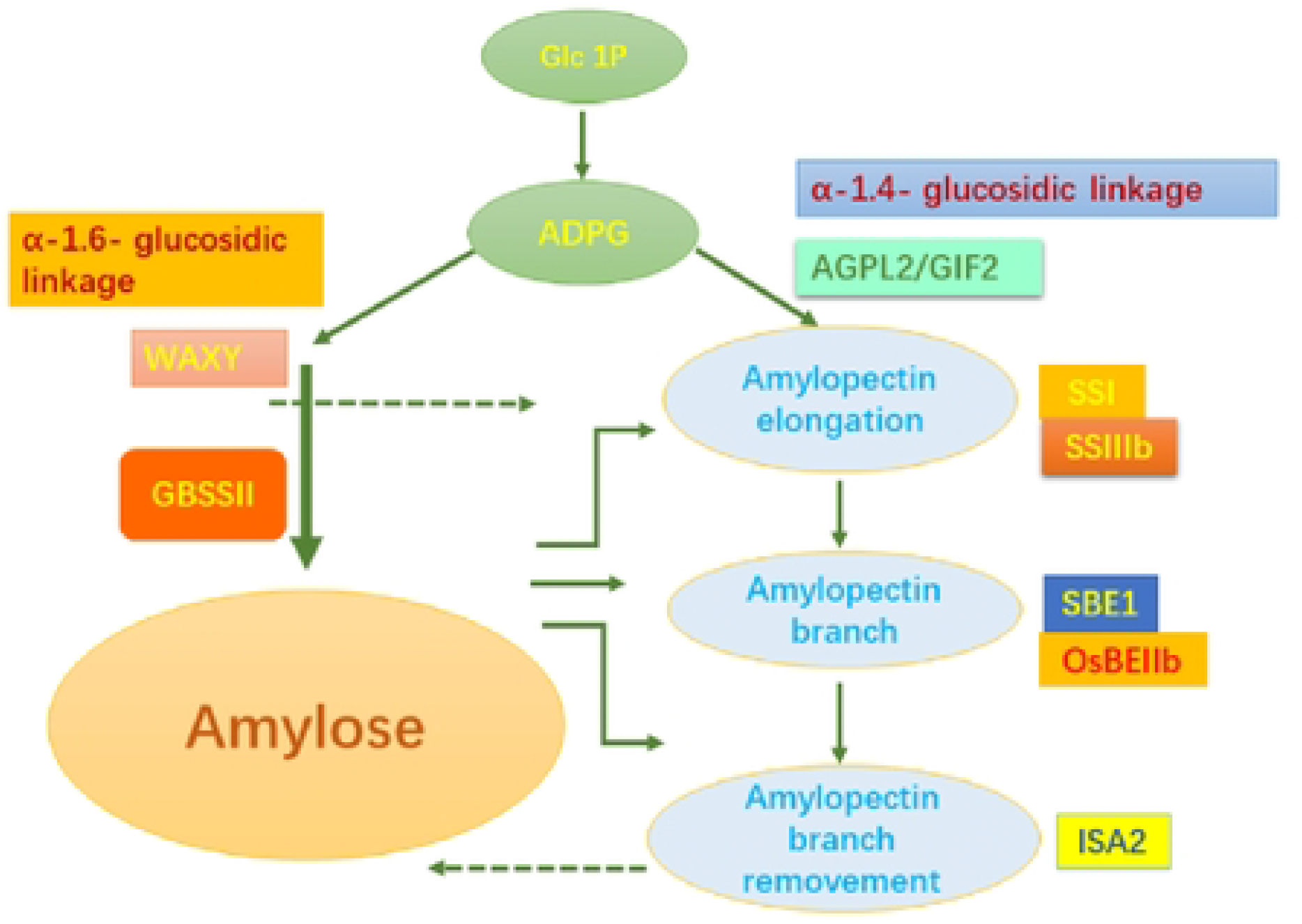
The model of starch synthesis in amyloplast/chloroplast of N2 mut. The pro-pro co-network during the early endosperm development in N2 mut, the crucial genes in starch synthesis are screened, which functions has been displayed clearly in the previous study.

**S5 Table.**
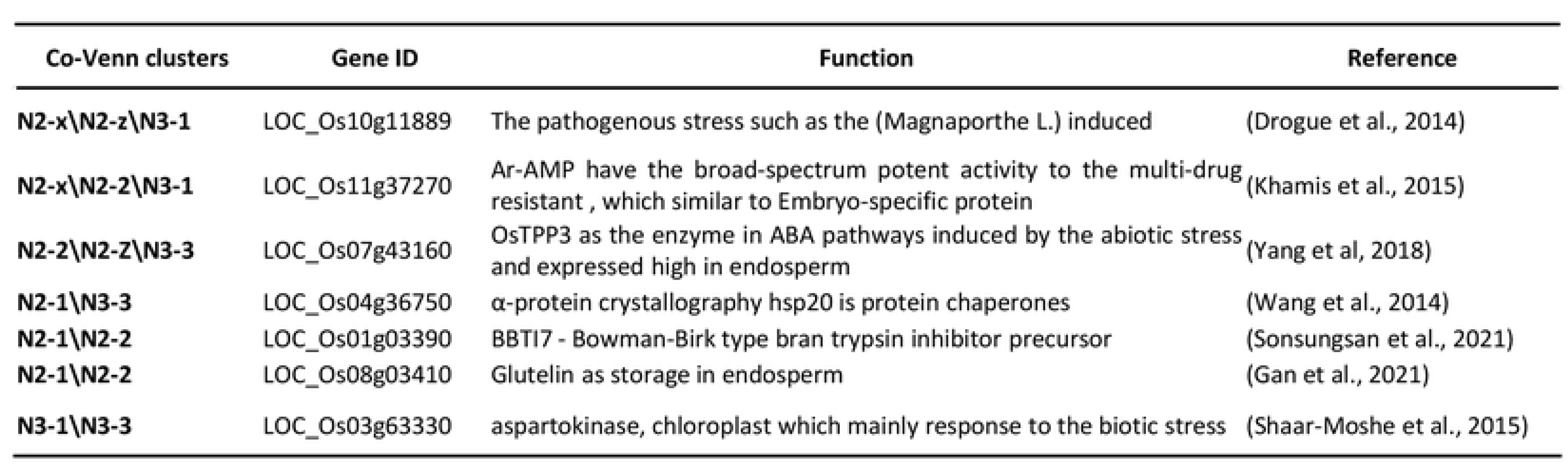
the genes co-expressed in the down-regulated Venn diagram (**N2 vs N3 vs WT, 4B Fig.**)

**S8 Table.** the volcano diagram of genes hub in DEGs of the seeds(**N2 vs N3 vs WT, 4B Fig.**)

| Up-regulated | All DEGs in top 5 | Function |
| --- | --- | --- |
| 1 | LOC_Os02g08150 | CCT/B-box zinc finger protein, putative, expressed |
| 2 | LOC_Os12g08810 | VTC2, putative |
| 3 | LOC_Os01g62810 | regulator of chromosome condensation, |
| 4 | LOC_Os04g09430 | cytochrome P450, |
| 5 | LOC_Os09g29210 | OsPUP2 |
| Down-regulated | LOC_Os01g01470 | OsNAC20; ONAC020 |
| 2 | LOC_Os02g15070 | glutelin, putative, expressed |
| 3 | LOC_Os11g13980 | hsp20/alpha crystallin family protein, putative, expressed |

**S10 Table.** the volcano plot of labeled genes in Sucrose and Starch pathway of the seeds (**N2 vs N3 vs WT, 4D Fig.**)

| Up-regulated | DEGs in the top 5 | Gene name | In Sucrose and Starch pathway |
| --- | --- | --- | --- |
| 1 | LOC_Os10g32810 | OsBAM2 | (Kim et al., 2017) |
| 2 | LOC_Os01g12020 | LTPL18 - Protease | (Zhang et al., 2010) |
| 3 | LOC_Os01g71670 | OsGLN2 | (Akiyama et al., 2004) |
| 4 | LOC_Os04g53310 | OsSSIIIb | (Wang et al., 2015a) |
| 5 | LOC_Os07g22930 | GBSSII | (Maung et al., 2021) |
| Down-regulated | LOC_Os06g49970 | alpha-amylase | (Damaris et al., 2019) |
| 2 | LOC_Os03g11420 | Os3BGlu6, glycosyl hydrolase family 1 | (Sun et al., 2021) |
| 3 | LOC_Os04g40510 | glycosyl hydrolase family 5 | (Xiang et al., 2019) |
| 4 | LOC_Os03g22790 | beta-amylase, | (Wang et al., 2022a) |
| 5 | LOC_Os10g40550 | CPuORF23 | (Shaar-Moshe et al., 2015) |

